# LPOR and the membranes – evolutionary pathway towards prolamellar body formation

**DOI:** 10.1101/2024.03.08.584095

**Authors:** Wiktoria Ogrodzińska, Katarzyna Szafran, Mateusz Łuszczyński, Olga Woźnicka, Michał Gabruk

## Abstract

Light-dependent protochlorophyllide oxidoreductase (LPOR) has captivated the interest of the research community for decades. One reason is the photocatalytic nature of the reaction catalyzed by the enzyme, and the other is the involvement of LPOR in the formation of a paracrystalline lattice called a prolamellar body (PLB) that disintegrates upon illumination, initiating a process of photosynthetic membrane formation.

In this paper, we have integrated three traditional methods previously employed to study the properties of the enzyme to investigate how LPOR evolved and how PLB forms. We found that in cyanobacteria, LPOR activity appears to be independent of lipids, with membrane interaction primarily affecting the enzyme post-reaction, with MGDG and PG having opposite effects on SynPOR. In contrast, plant isoforms exhibit sequence alterations, rendering the enzyme effective in substrate binding mainly in the presence of anionic lipids, depending on residues at positions 122, 312, and 318. Moreover, we demonstrated that the interaction with MGDG could initially serve as enhancement of the substrate specificity towards monovinyl-protochlorophyllide (Pchlide). We have shown that the second LPOR isoforms of eudicots and monocots accumulated mutations that made these variants less and more dependent on anionic lipids, respectively. Finally, we have shown that in the presence of Pchlide, NADP+, and the lipids, plant but not cyanobacterial LPOR homolog remodel membranes into the cubic phase. The cubic phase is preserved if samples supplemented with NADP+ are enriched with NADPH.

The results are discussed in the evolutionary context, and the model of PLB formation is presented.

**Significance:** LPOR is a unique enzyme with photocatalytic properties, developed by cyanobacteria and inherited by algae and plants. In this study, we investigated the properties of the cyanobacterial homolog, revealing that two lipids, PG and MGDG, have opposite effects on enzyme activity. Additionally, we identified mutations in plant isoforms that render the enzyme dependent on anionic lipids. Moreover, we demonstrated that in the presence of NADP+, the plant homolog remodels lipids into a cubic phase, which appears to be the initial step of prolamellar body (PLB) formation. PLB is a unique paracrystalline arrangement of lipids and proteins found in immature chloroplasts, which disintegrates upon illumination, initiating photosynthetic membrane formation.

## Introduction

Chlorophyll molecules have been essential elements of the photosynthetic machinery for billions of years. During this time, phototrophic organisms have not only improved the efficiency of the light capturing reaction^1^ and carbon assimilation^2,3^ but have also perfected the enzymes involved in chlorophyll biosynthesis (Video S1) and their regulatory network. Around 1.4 billion years ago (GYA), cyanobacteria developed an alternative enzyme to catalyze the penultimate reaction of the pathway^4,5^—specifically, the reduction of protochlorophyllide (Pchlide)—in an oxygen-insensitive manner, utilizing light as a trigger for the reaction. The enzyme, known as light-dependent protochlorophyllide oxidoreductase (LPOR), uses the excited state of Pchlide trapped in the pigment binding pocket to reduce one of the double bonds of protochlorophyllide (Pchlide), with the aid of NADPH^6^. The exact molecular mechanism of the photocatalytic reaction is still under debate, however, it is widely agreed that the ternary complex of Pchlide:LPOR:NADPH produces NADP^+^ and chlorophyllide (Chlide) as a result of the reaction.

LPOR has captured the interest of the research community for decades, making it one of the most studied enzymes of the pathway^7^ for at least two reasons. One is the light-dependent nature of the reaction that plays a regulatory role in both chloroplast maturation and photomorphogenesis^7,8^. The other reason is how angiosperm plants utilize properties of LPOR to prepare for photosynthetic activity when light is limited, for example, during seedling germination or within leaf buds. During prolonged periods of darkness, immature chloroplasts, or etioplasts, accumulate lipids and the ternary LPOR complexes in a form of a paracrystalline three-dimensional lattice called a prolamellar body (PLB) that resembles a cubic phase^9^. PLBs are stable in darkness, but as soon as exposed to light, LPOR catalyzes its reaction, and the paracrystalline structure of PLB starts to disintegrate, initiating a process of photosynthetic membrane formation. These complexes, therefore, serve as a reservoir for lipids and chlorophyll intermediates for developing photosynthetic membranes, allowing for a quicker assembly of the photosynthetic apparatus and consequently increased photoautotrophic capacity.

However, the evolutionary origin of PLB is unknown, as well as the molecular mechanism of its formation. A few premises suggest that the interaction between LPOR and lipids is involved. Firstly, monogalactodiacylglycerol (MGDG), the main lipid component of PLB, can obtain a cubic phase, but not under physiological conditions^10^. Secondly, two pools of Pchlide can be distinguished within PLB: one not bound to enzymatically active complexes and the other undergoing photoconversion upon illumination. The latter has a distinguished emission maximum of fluorescence at 655 nm, when measured at 77K^11^. Experiments using recombinant plant LPOR variants revealed that to reconstruct complexes with such emission maximum, the reaction mixture must be supplemented with MGDG^12^. This observation indicates that LPOR interacts with MGDG within PLB. Beside MGDG, also anionic lipids^12^, such as phosphatidylglycerol (PG), were shown to be important for effective formation of active LPOR complexes, with both lipids playing different roles^12^.

Anionic lipids have been shown to increase the affinity of plant LPOR variants to NADPH^12^, while MGDG was shown to trigger the oligomerization of the enzyme^13^. Structural analysis of these oligomers, using cryoelectron microscopy, revealed that LPOR subunits form lines of dimers that wrap around thin tubes of lipid bilayer, producing long, linear filaments of helical nature in vitro^13^. Within these complexes, each LPOR subunit binds NADPH and the pigment, partially embedded in the outer leaflet of the bilayer. Surprisingly, a galactosyl group of MGDG is bound next to the Pchlide within the pigment binding pocket, which consists of two parts. One is an alpha helix motif, named helix α10, and the other is a loop called Pchlide loop^13^. Both structural motifs sandwiching the pigment and trap it within the enzyme in the catalytic conformation, where a photon excites Pchlide to trigger the reaction^13^.

A cryoelectron tomography study of isolated PLBs confirmed the presence of helical assemblies of LPOR within the structure^14^ but did not explain how its cubic phase emerges. It was suggested that other proteins could be responsible for the emergence and maintenance of a cubic phase of PLB, namely CURT1 proteins since the absence of CURT1 proteins has been shown to severely perturbs PLB organization^15^. Additionally, it is unclear if cyanobacterial isoform of LPOR can produce PLB. The variant from Synechocystis (SynPOR) does not form complexes having emission maximum of 655 nm in a presence of MGDG, unlike plant isoforms^4^. On the other hand, heterologous expression of cyanobacterial LPOR was shown to compensate for the reduced size of PLB in AtPORA knock-out plants^16^. PLBs have been reported in some algae species^17^, which have one copy of LPOR gene. In vascular plants, the number of copies of the LPOR gene typically ranges from one to three.

It has been shown that ancestors of monocots and eudicots underwent separate duplication events, leading to the independent emergence of two isoforms separately in monocots and eudicots^4^. In the latter, also secondary LPOR duplications and gene deletions have been identified^4^, leading, for example, to the emergence of three isoforms of *A. thaliana*, named AtPORA, B and C. The role of these distinctive LPOR isoforms is unknown, but they are differently regulated^6^, what indicates that they may have different biochemical properties regarding substrate binding or allosteric regulation. Even though the protein identity of plant LPOR sequences is high^4,5^, a handful of mutations could lead to distinctive properties. So far, differences in the affinity towards NADPH have been reported for three isoforms of Arabidopsis, but without any molecular explanation^4^.

In this paper we wanted to characterize the influence of the lipids, namely MGDG and PG, on the activity of cyanobacterial LPOR variants. Additionally, we investigated how these interactions were modified in plant variants of the enzyme. Results presented in the paper shed a new light on evolution of LPOR and provide a model of the PLB formation in plants.

## Results

### Dinucleotide binding in SynPOR and lipids

To determine the effect of lipids on bacterial isoform of LPOR, we used an enzyme from Synechocystis (hereafter SynPOR, Uniprot: Q59987) as a model protein. Firstly, we analyzed the shifts of the fluorescence emission maxima of complexes with two pigments: Pchlide, the substrate, and Chlide, the product of the reaction that is generated during illumination, added directly to the reaction mixture.

We noticed that the sole presence of SynPOR in the reaction mixture red shifts the emission maximum of both pigments by 4-5 nm (Fig. 1AC). Introduction of NADP^+^ into the lipid-free mixture induced further red shift in the emission maximum of the complex with Pchlide by 7 nm, but not with Chlide. For the latter, the effect of NADP^+^ was only observed if the reaction mixture was supplemented with PG – in such case the Chlide:LPOR:NADP^+^:PG complex had an emission maximum at 686 nm (Fig 1C). The presence of NADPH influenced the complexes with both Pchlide and Chlide, red shifting the emission maxima to 647 and 689 nm, respectively, in the lipid-free mixture. The addition of PG to these complexes did not affect the emission maximum of the complex with Pchlide, but red-shifted the emission of the complex with Chlide by additional 3 nm. The measurements allowed us to assign specific emission maxima to the complexes of different composition (Fig. 1AC). We then utilized these distinctions in the emission maxima to interpret the spectra of the illuminated reaction mixtures where Chlide is generated.

**Figure 1.**
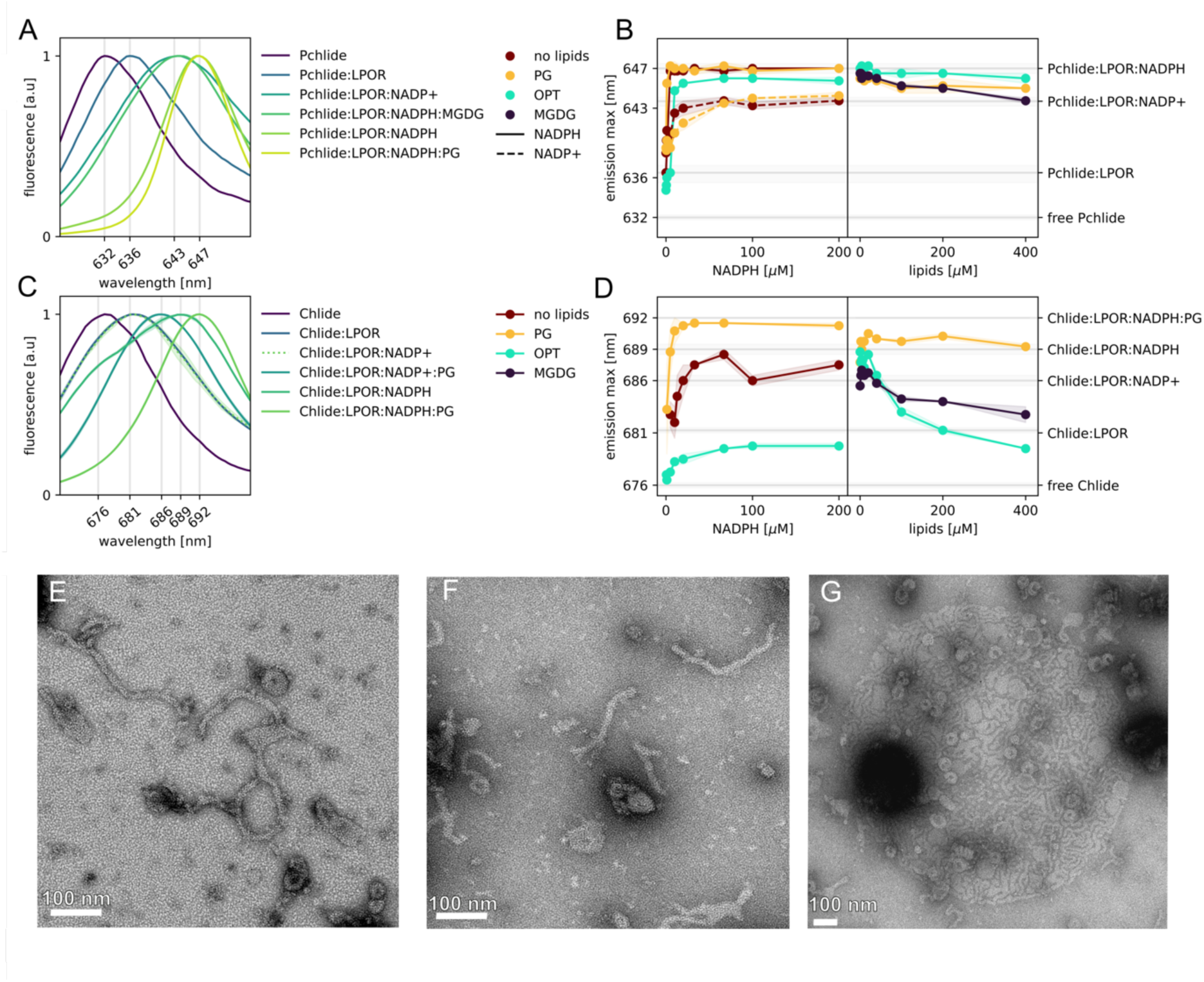
Cyanobacterial SynPOR and lipids. **AC.** The fluorescence emission spectra of Pchlide (**A**) and Chlide (**C**) in reaction mixtures of different composition. **BD.** The position of the emission maximum of Pchlide before illumination (**B**) and Chlide generated after 20 seconds of illumination (**D**) for SynPOR in different NADPH and lipids concentrations. The reaction mixture contained 5 µM Pchlide, 15 µM SynPOR, 40 µM lipids (left panels), 200 µM dinucleotide (right panels). Solid and dotted lines represent mixtures supplemented with NADPH or NADP+, respectively. All spectra used for this figure are presented in **Fig. S1**. **EFG.** Negative-stain electron microscopy micrographs of samples supplemented with PG and Chlide:SynPOR:NADPH (**E**) and Pchlide:SynPOR:NADP+ (**FG**).

We found that in the lipid-free environment and at low NADPH concentrations the emission maximum of the reaction mixture resembles that of Chlide:LPOR complex (Fig. 1D). The more NADPH, the emission maximum shifts, firstly to the one resembling Chlide:LPOR:NADP^+^, and then continuously shifts towards the emission characteristic for the Chlide:LPOR:NADPH complex. In the presence of PG, the emission maximum also depends on the concentration of NADPH, but in general it is much more red-shifted than for the complexes formed in the lipid-free environment even for low PG concentrations (Fig. 1D). The addition of a lipid mixture which has been previously utilized in the cryoEM study of plant AtPORB and referred to as the OPT lipids (50 mol% MGDG, 35 mol% DGDG, and 15 mol% PG), cause in general a blue-shift of the emission maximum compared to the complexes that are formed in the lipid-free environment. The emission maximum of Chlide that formed in the reaction mixture containing OPT lipids resembled the one determined for free Chlide or Chlide:LPOR complex, depending on NADPH and OPT concentrations. A similar effect was observed with the addition of MGDG: the higher the concentration of MGDG, the greater the blue shift in the emission maximum, towards the emission characteristic to Chlide:LPOR complexes.

Surprisingly, the lipids did not affect the formation of the complexes with Pchlide, except the observation that increasing lipids concentrations (either PG, OPT or MGDG) result in enhanced blue-shift of the emission maximum (Fig. 1B). The maximum shifted from 647 nm in the lipid-free environment to 643 nm for high MGDG concentrations, which is comparable to emission determined for the Pchlide:LPOR:NADP+ complex (Fig. 1A). However, these complexes do not contain NADP^+^, because these reaction mixtures were supplemented with NADPH and the complexes were enzymatically active (Fig. S1H). These examples provide yet another piece of evidence that the relationship between the microenvironment of Pchlide/Chlide within the LPOR’s binding pocket and the emission maximum of the complex is a complicated issue, since two different complexes can emit light of the same wavelength, at least at the resolution of our spectrophotometer.

Additionally, we investigated the presence of large oligomers in all of the investigated conditions with the use of electron microscopy. We detected such complexes only for the samples supplemented with PG. A small number of thin bended filamentous complexes were found in the samples containing SynPOR, Chlide, NADPH and PG (Fig. 1E). Similar complexes were much more abundant when samples with SynPOR and PG were supplemented with NADP^+^ and Pchlide (Fig. 1F). In these conditions, the complexes were also observed to condense into larger structures (Fig. 1G). We conclude that PG enhances dinucleotide binding and oligomerization of SynPOR, whereas MGDG primarily influences enzyme post-reaction, promoting products release.

### Role of helix α10 in Pchlide binding

Subsequently, we wanted to investigate the role of two parts of the enzyme responsible for Pchlide binding: the Pchlide loop and helix α10 (hereafter α10) (Fig S2, Fig. 2AB). We generated hybrid mutants of AtPORB, in which part of the plant sequence, either the loop or the helix, has been substituted with corresponding fragments from SynPOR. Out of these two hybrid proteins, AtPORB-Pchlide_loop has not been enzymatically active, while AtPORB_α10 could be thoroughly characterized (Fig. S2, Fig. 2).

**Figure 2.**
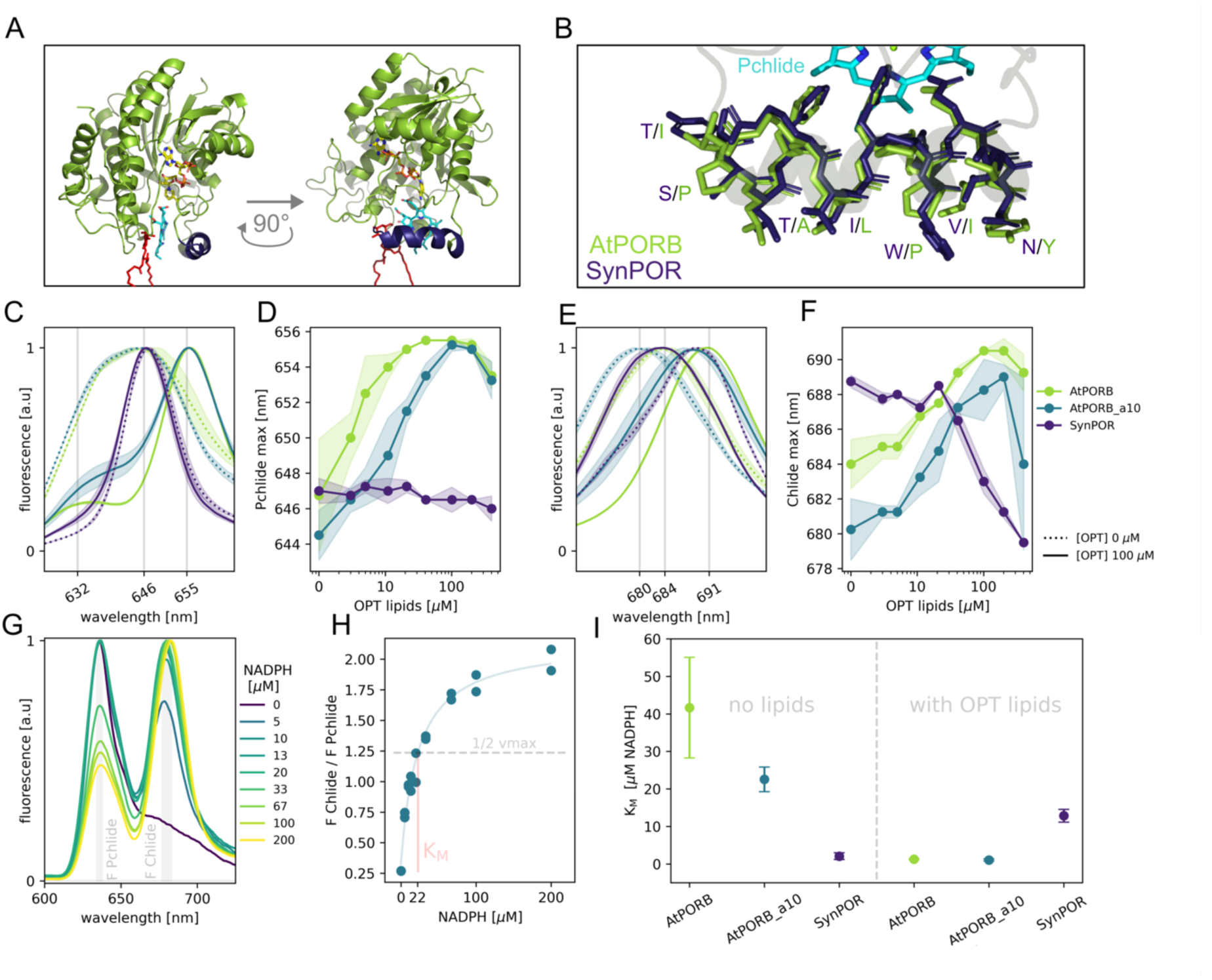
Role of helix α10. **A.** The position of helix α10 (dark purple) within AtPORB (green). MGDG (red) and Pchlide (cyan) are shown. **B.** The residues of helix α10. Residues that differ between AtPORB (green) and SynPOR (dark purple) are labeled. **CE.** The fluorescence emission spectra of reaction mixtures containing 15 µM LPOR (either AtPORB, SynPOR, or hybrid AtPORB_α10), 5 µM Pchlide, 200 µM NADPH without (dotted lines) and with (solid lines) 100 µM OPT lipids before (C) and after 20 seconds of illumination (E). Shared areas represent the standard deviation between replicates. **DF.** The relationship between OPT lipid concentration and the position of the maximum fluorescence emission of Pchlide before illumination (D) and Chlide after 20 seconds of illumination (F). The composition of the reaction mixtures is as in CE, except for the OPT lipids. Shared areas represent the standard deviation between replicates. **G.** Exemplary spectra of illuminated reaction mixtures of hybrid AtPORB_α10. The relative intensities of remaining substrate (F Pchlide) and generated product (F Chlide) were read from each spectrum, and a ratio of F Chlide / F Pchlide was calculated for K_M_ determination. **H.** The relationship between F Chlide / F Pchlide and NADPH concentration for hybrid AtPORB_α10 in the absence of lipids. A fit of a modified Michaelis-Menten equation is shown, and the graphical representation of K_M_ and ½ V_max_ is shown. **I.** Values of K_M_ determined for AtPORB, SynPOR, and hybrid AtPORB_α10 in the absence and presence of 40 µM OPT lipids. Error bars represent the uncertainty associated with the fitted constant K_M_. All spectra used to prepare this figure are presented in Figure S3. The fits of a modified Michaelis-Menten equation are presented in Figure S4. The exact values of the determined K_M_ constants are presented in Table S1.

The structural comparison of AtPORB with SynPOR (the Alphafold2 prediction downloaded from Uniprot database^18^) revealed that the residues of the helix α10 interacting with Pchlide are identical in both proteins, but seven of the residues are different (Fig. 2AB). We found that these residues are not involved in the formation of the complexes having an emission maximum at 655 nm when OPT lipids are present in the reaction mixture, since the spectrum of a hybrid AtPORB_α10 mutant resembled that obtained for AtPORB_WT (Fig. 2CD). However, higher lipid concentrations were required for the mutant than for the WT enzyme to produce complexes having similar emission maxima (Fig. 3D). Moreover, the blue shift of Chlide emission maximum in a presence of MGDG or OPT lipids, observed for SynPOR after the exposure of the reaction mixture to light (as depicted in Fig. 1D), was not observed for the AtPORB_α10 mutant (Fig. 3EF).

**Figure 3.**
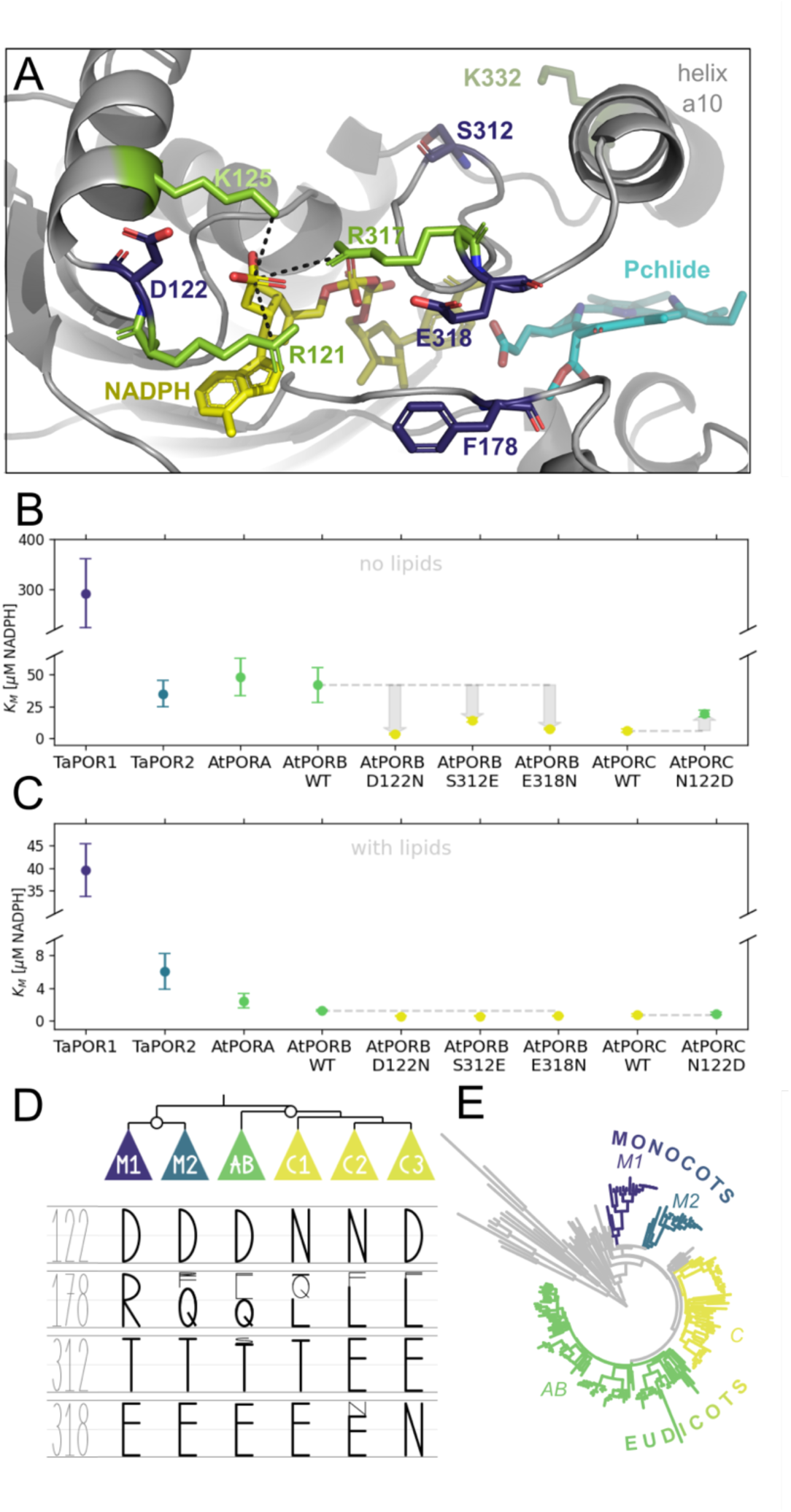
Residues involved in NADPH binding in LPOR. **A.** Part of the AtPORB structure showing residues involved in NADPH binding. NADPH (yellow) and Pchlide (cyan) are shown, and helix α10 is labeled. **BC.** Values of the KM constant determined for two isoforms of wheat (TaPOR1, TaPOR2), three isoforms of A. thaliana (AtPORA, AtPORB, and AtPORC), and mutants of AtPORB and AtPORC in the absence (B) and presence of 40 µM OPT lipids (C). Reaction mixtures contained 15 µM LPOR, 5 µM Pchlide, and NADPH. Error bars represent the uncertainty associated with the fitted constant K_M_. **D.** A simplified cladogram of LPORs of angiosperms showing the conservation of residues at positions 122, 178, 312, and 318 (numbering for AtPORB). The size of letters is scaled proportionally to the number of sequences that have a given amino acid at the given position. Subclades of clade C are distinguished. **E.** The phylogenetic tree of eukaryotic LPORs. Two clades of variants of monocots (M1 and M2) and two clades of variants of eudicots (AB and C) are colored. All spectra used to prepare this figure are presented in Figure S5. The fits of a modified Michaelis-Menten equation are presented in Figure S6. The exact values of K_M_ constant are presented in Table S1.

To investigate the role of helix α10 in the active complex formation, we used a quantitative assessment of NADPH binding, namely an analogue of Michaelis-Menten constant (K_M_) using samples with increasing NADPH concentrations illuminated for 20 seconds (Fig. 2GH, Materials and Methods).

The K_M_ constant was the highest for AtPORB WT when no lipids were present, but it dropped by nearly two orders of magnitude when the OPT lipids were added to the mixture (Fig. 2I, Table S1). For SynPOR it was the contrary, the K_M_ constant was low when no lipids were present, and it increased when the OPT lipids were added to the mixture. For a hybrid AtPORB_α10, an improved NAPDH binding was observed in comparison to AtPORB WT in lipid-free reaction mixture, and a significant drop of K_M_ constant was noticed after the addition of the lipids, similarly to AtPORB WT (Fig. 2I, Table S1).

In order to examine which other residues, beside helix α10, can influence the active complex formation in LPOR variants of plants, we performed site-directed mutagenesis study inspired by natural variations found in an LPOR sequence dataset, published previously^4^. We noticed that all eukaryotic LPOR sequences have conserved positively charged residues interacting with the phosphate group of NADPH at positions 121, 125 and 317 (numeration according to AtPORB sequence, Fig. S7).

Interestingly, in close proximity to these positions, we found residues that are differently conserved within the clades of angiosperms, namely at positions 122, 312 and 318 (Fig. 3DE). Two clades of monocots (hereafter: M1 and M2, that TaPOR1 and TaPOR2 from *T. aestivum* belong to, respectively) and one clade of eudicots (hereafter: clade AB, that AtPORA and AtPORB belong to) have conserved D, T (or S in the case of 9.6% of all of the sequences) and E, respectively. Interestingly, the other clades of eudicots (hereafter: clade C1-3, that AtPORC belong to) have D, E and N at these positions, in different combinations (Fig. 3DE). Therefore, we split the clade C into three subclades, depending on the residues at positions 122 and 318 (Fig. 3D, supplementary dataset 1).

To determine the role of these substitutions on NADPH binding we performed site-directed mutagenesis using two LPOR variants, AtPORB and AtPORC, which belong to the clades AB and C1, respectively (Fig. 3BC). Again, we employed the K_M_ constant, an analogue of Michaelis-Menten constant, to quantitively assess the NADPH binding. We noticed that substitutions D122N, S313E and E318N decreased the values of K_M_ for AtPORB when no lipids were present in the reaction mixture. At the same time, the N122D substitution for AtPORC increased the value of K_M_ under the same conditions (Fig. 3C). Among the analyzed WT variants in the absence of lipids, TaPOR2, AtPORA, and AtPORB exhibited similar values for the K_M_ constant, whereas TaPOR1 displayed notably higher values, and AtPORC showed significantly lower values (Fig. 3B). The presence of lipids lowered K_M_ to similar values for all analyzed variants, except for TaPOR1 (Fig. 3C). For the latter, the presence of lipids also lowered K_M_, but to a higher value than for the rest of the analyzed enzymes.

### MGDG and Pchlide binding

In the cryoEM structure of AtPORB, we found that MGDG molecule can interact with two parts of Pchlide in the binding pocket of the enzyme due to the close proximity^19^: with the carboxyl at C13^1^ and with a sidechain at C8. The C8 sidechain is especially interesting because two forms of Pchlide exist within the Chl biosynthesis pathway that differ at this position. One, MV-Pchlide, has an ethyl group at this position, while the other, DV-Pchlide, a vinyl group (Fig. 4D). To investigate whether the interaction between MGDG and sidechain at C8 of Pchlide affects the formation of active complexes in LPOR variants from plants and cyanobacteria, we used TaPOR2 (isoform 2 of T*. aestivum*, member of the M2 clade (Fig. 4DE)) and SynPOR.

**Figure 4.**
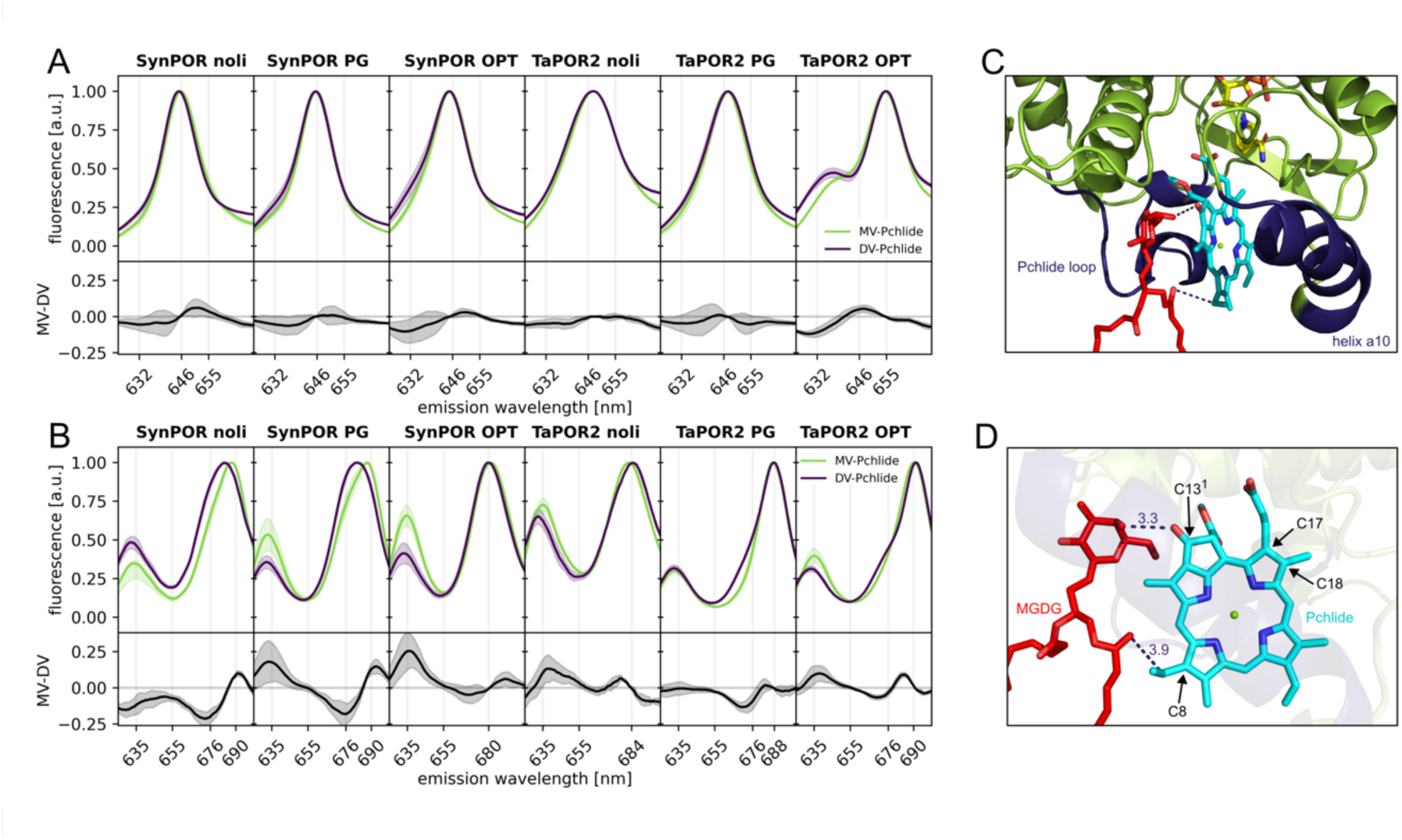
Group at C8 of Pchlide and the influence of MGDG on LPOR. **AB.** The spectra of the reaction mixtures containing SynPOR or TaPOR2 (from *T. aestivum)* with MV-Pchlide (green) and DV-Pchlide (dark purple) before (A) and after 20 minutes of illumination (B). The reaction mixtures contained 15 µM LPOR, 3.2 µM Pchlide (either MV- or DV-), 200 µM NADPH, and either 100 µM PG, 100 µM OPT, or no lipids (Noli). Shared areas represent the standard deviation between replicates. Corresponding difference spectra (MV minus DV) are presented at the bottom of panels A and B, for which the shared areas represent the overall standard deviation between replicates obtained for MV- and DV-Pchlide. **CD.** Position of MGDG (red) and Pchlide (cyan) within LPOR, sandwiched between the Pchlide loop and helix α10 (dark purple). MV- and DV-Pchlide differ with a group at C8 of Pchlide. Selected carbon atoms of Pchlide are labeled. Distances between MGDG and Pchlide are in Å. All spectra used to prepare this figure are presented in Figure S8.

We noticed that under the examined conditions neither SynPOR nor TaPOR2 can discriminate between MV- and DV-Pchlide when the reaction mixture is either supplemented with PG or no lipids are added: the emission maxima for both Pchlide forms are nearly identical and the difference spectra are flat within the uncertainty (Fig 4). However, when OPT lipids were added to the reaction mixture, the spectra of samples with TaPOR2, but not SynPOR, exhibited differences between complexes with MV- and DV-Pchlide. TaPOR2 samples with MV-Pchlide in the presence of OPT lipids had a smaller shoulder at 632 nm than samples with DV-Pchlide. This suggests a higher affinity of TaPOR2 in the presence of OPT lipids towards MV-Pchlide compared to DV-Pchlide (Fig. 4A).

Furthermore, it was observed that samples containing MV- and DV-Pchlide exhibited variations in the emission maximum of newly generated Chlide. These differences were minor after 20 seconds of illumination (Fig. S8) but became more pronounced with longer exposure. After 20 minutes of illumination, TaPOR2 showed some differences in the emission spectra of reaction mixtures between newly formed MV- and DV-Chlide, namely a higher shoulder at 676 nm for DV-relative to MV-Chlide if any lipids were present in the reaction mixture (Fig. 4B). For SynPOR, however, the emission maxima of DV-Chlide were substantially blue-shifted in comparison to those of MV-Chlide in absence of lipids or in sole presence of PG. The emission maxima of both MV- and DV-Chlide were always at 680 nm when SynPOR mixtures were supplemented with OPT lipid mixture. Additionally, less MV-than DV-Pchlide were converted into Chlide when the reaction mixtures of SynPOR were supplemented with any lipids (Fig. 4B). These results demonstrate that MV- and DV-Pchlide are indistinguishable for LPOR isoforms, unless there are lipids containing MGDG. Only then MV-Pchlide is preferentially sequester in LPOR oligomers of plant isoforms. However, there are differences in the disassembly of the complexes with MV- and DV-Chlide, generated in the photocatalytic reaction, for both cyanobacterial and plant isoform.

### NADP(H) and the oligomerization of plant LPOR

Then, we investigated NADP^+^ binding to AtPORB, as a representative plant isoform. We noticed that AtPORB:Pchlide can bind NADP^+^ in the presence of OPT lipids and the resulting complex has an emission maximum at 647 nm, while the complexes with NADPH have maximum at 655 nm (Fig. 5A). Importantly, the emission maximum of the reaction mixture supplemented with NADP^+^ and the lipids resembles the one of the active ternary LPOR complex in lipid-free environment. However, the complexes with NADP+ were not enzymatically active (Fig. S9E): no Chlide fluorescence was detected in samples supplemented solely with NADP+ after exposure to light. For complexes with each of the dinucleotides, the emission maximum of Pchlide depends on the lipid concentration (Fig. 5B). The higher the lipids concentration, the more red-shifted the maximum. But when the OPT concentration increased above 100 μM, a blue shift of the emission maximum can be seen (Fig. 5B). While investigating the relative fluorescence intensities of the ternary complexes with NADP+ and NADPH, with a maximal emission at 647 and 655 nm, respectively, in the range of the OPT lipids concentrations, we noticed a similar shapes of both curves (Fig. 5C), with a clearly visible maximum at 100-200 μM OPT.

**Figure 5.**
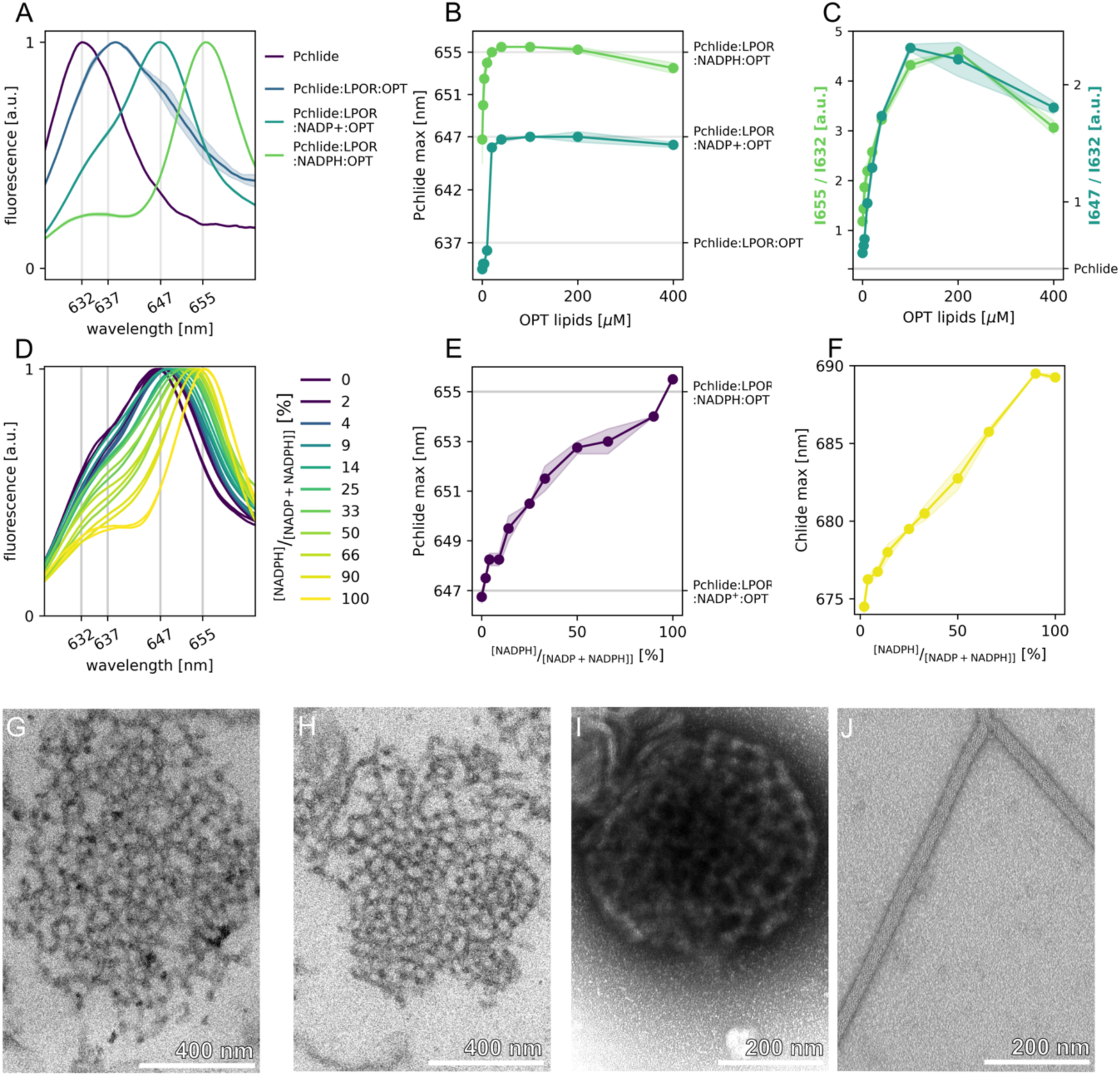
NADP+ binding to Pchlide:LPOR complexes in the presence of lipids results in a cubic phase. **A.** The fluorescence emission spectra of different Pchlide complexes. The reaction mixtures contained: 5 µM Pchlide, 15 µM AtPORB, 200 µM NADP(H), and 100 µM OPT. Shared areas represent the standard deviation between replicates. **B.** The relationship between OPT lipid concentration and the position of the maximum fluorescence emission of 5 µM Pchlide before illumination in the presence of NADPH or NADP+ (200 µM) and AtPORB (15 µM). Shared areas represent the standard deviation between replicates. **C.** The relationship between OPT lipid concentration and the relative fluorescence intensity of complexes with NADPH (655/632, green series, left axis) or with NADP+ (647/632, teal series, right axis). A grey horizontal line labeled “Pchlide” represents the 655/632 and 647/632 ratio for pure Pchlide in the buffer. Shared areas represent the standard deviation between replicates. **D.** The fluorescence emission spectra of Pchlide (5 µM) in the presence of AtPORB (15 µM) and OPT lipids (40 µM) with different contents of NADPH and NADP+ (0-400 µM). Exact concentrations are provided in Fig. S9DE. **EF.** The relationship between OPT lipid concentration and the position of the fluorescence emission of: Pchlide before illumination (E); and newly generated Chlide after 20 seconds of illumination (F). The composition of the reaction mixtures is as in D. Shared areas represent the standard deviation between replicates. **GH.** Transmission electron microscopy micrographs of cross-sections of a reaction mixture containing Pchlide (5 µM), AtPORB (15 µM), OPT lipids (100 µM), and NADP+ (200 µM) (G); and a reaction mixture as in G to which NADPH (200 µM) was added after initial incubation (H). **IJ.** Negative-stain transmission electron microscopy micrographs of reaction mixtures containing Pchlide (5 µM), AtPORB (15 µM), OPT lipids (100 µM), and dinucleotide (200 µM): either NADP+ (I) or NADPH (J).

Then, we investigated the presence of LPOR oligomers in samples with NADP^+^ and NADPH using negative staining and electron microscopy. The AtPORB samples supplemented with Pchlide, OPT lipids and NADPH contain long, linear, filamentous oligomers, as we previously described and characterized^13^ (Fig. 5J). However, the samples supplemented with Pchlide, OPT lipids and NADP^+^ contained spherical assemblies of branching tubes (Fig. 5I). To determine the inner structure of these assemblies we prepared identical samples, containing NADP^+^, but used different staining technique that involves sample embedding in the resin and a subsequent microsectioning with the microtome. This approach revealed that complexes of LPOR, Pchlide, OPT lipids and NADP^+^ resemble a cubic phase, although they are not that symmetric like PLB (Fig. 5G). This implies that the formation of the characteristic paracrystalline lattice of PLB requires some additional factors or steps.

Then we analyzed the effect of subsequent supplementation of a reaction mixture containing Pchlide, LPOR, NADP^+^ and OPT lipids with NADPH in darkness. The more enriched with NADPH the reaction mixture is in darkness, the more red-shifted is the emission maximum of Pchlide, which shifts from 647 nm for the sample with just NADP^+^, up to approximately 655 nm for the sample containing supplemented with just NADPH (Fig. 5DE). The presence of NADPH resulted in the formation of enzymatically active complexes (Fig. S9E), even though it was NADP^+^ that was added first, what suggests that NADPH can displace NADP^+^ in the binding pocket. Moreover, the more NADPH in comparison to NADP^+^ added to the reaction mixture, the higher relative fluorescence intensity of Chlide (Fig. S9E) and more red-shifted its emission maximum (Fig. 5F). Intriguingly, the cubic phase of LPOR complexes was preserved when NADPH was added to the reaction mixture after initial incubation with NADP^+^ (Fig. 5H). This suggests that the order of dinucleotide binding to the enzyme determines the shape of LPOR oligomers formed on lipid membranes.

## Discussion

LPOR is the most extensively studied enzyme in chlorophyll biosynthesis with a rich historical record of research and investigation^6,11^. In this paper, we have integrated three traditional methods that have been previously employed to study the properties of the enzyme. Low-temperature fluorescence measurements offer insight into the core of the reaction, where the excited state of the pigment plays a pivotal role. The sensitivity of the shifts of the fluorescence emission maxima allow to monitor the changes in the pigment microenvironment, so to track Pchlide binding, oligomerization, and product release^20–22^. Electron microscopy enables visualization of membrane shapes influenced by interaction with the enzyme ^23,24^. Finally, the use of recombinant proteins and purified pigments provides an opportunity to investigate in vitro the enzyme’s properties under different conditions in an isolated system, without interference from other proteins or molecules^12,25–27^.

This experimental approach, combined with the analysis of hundreds of sequences in an evolutionary context, published before^4^, enabled us to uncover the evolutionary history of LPOR and shed new light on the mechanism of PLB formation and PLB-emerging events.

### LPOR in cyanobacteria

Several previous studies, along with the data depicted in Fig. 1, suggest that SynPOR effectively binds substrates and facilitates the reaction in the absence of lipids ^30–32^. Interestingly, the presence of lipids appears to moderately disrupt the formation of the active complex, as evidenced by an increase in the K_M_ constant for NADPH binding (Fig. 2).

Surprisingly, interaction with MGDG appears to facilitate the release of the dinucleotide from the complex post-reaction. This is indicated by similar effects observed on the fluorescence emission maximum of newly generated Chlide with both OPT lipids and pure MGDG (Fig 1D). The release of the dinucleotide likely involves a conformational change in the enzyme, as NADP(H) is deeply sequestered within the protein scaffolding in its active conformation. MGDG likely assists in this conformational change by stabilizing the open apo conformation of cyanobacterial LPOR. That would also explain why the OPT lipids increase the value of K_M_ for NADPH binding to SynPOR.

The observations discussed above suggest that in *Synechocystis* cells, SynPOR primarily binds Pchlide and catalyzes reactions in a soluble form, however, NADP^+^ and Chlide release occur exclusively at membranes rich in MGDG (Fig. 6A), likely facilitating the transfer of the final metabolic intermediate of Chl synthesis to the membrane for the last catalytic step of the integral membrane protein - chlorophyll synthase.

**Figure 6.**
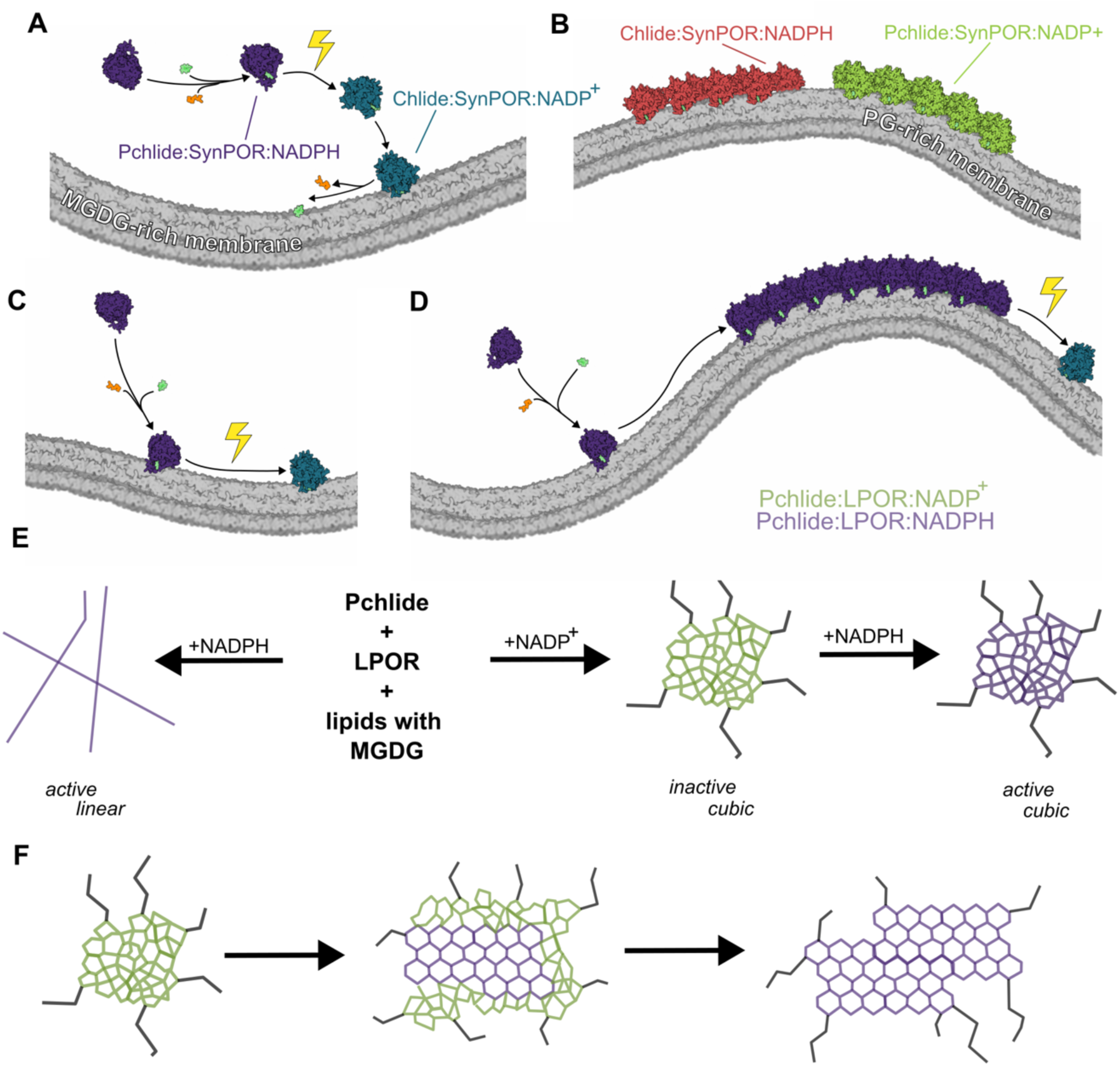
LPOR in cyanobacteria, plants, and hypothetical intermediate variants. **AB.** A model of SynPOR action. The formation of the active Pchlide:SynPOR:NADPH complex is most effective in solution, but an MGDG-rich membrane is required for the effective release of products generated after illumination (A). The PG-rich membrane promotes the formation of stable oligomers of Chlide:SynPOR:NADPH and Pchlide:SynPOR:NADP+ (B). **C.** A hypothetical intermediate variant of LPOR evolution, in which mutations close to the NADPH binding site made the ternary complex formation dependent on anionic lipids for prolonged storage of active complexes on the membrane, but oligomerization interfaces of the enzyme were not yet developed. **D.** A hypothetical intermediate variant of LPOR evolution, in which the ternary complexes on the membrane with anionic lipids developed oligomerization interfaces, leading to the formation of oligomers in darkness able to remodel membranes. **E.** A schematic representation of the oligomeric forms that modern-day LPOR variants of plants can form. The mixture of LPOR, Pchlide, and lipids containing MGDG can form linear active filaments when supplemented with NADPH, but produces a cubic inactive phase when supplemented with NADP+. The cubic phase is preserved, and activity restored if samples supplemented with NADP+ are enriched with NADPH after the initial incubation with the former. **F.** A model of prolamellar body formation in plants. Initially, the NADP+ concentration is high, allowing the formation of Pchlide:LPOR:NADP+ complexes on membranes with a cubic phase. Only then does the NADPH concentration slowly increase, displacing NADP+ within LPOR, and leading to the crystallization of Pchlide:LPOR:NADPH complexes and the formation of the characteristic paracrystalline lattice of PLB. Cartoons in A-C were created with Illustrate^28^ and CellPAINT^29^.

Moreover, our experiments revealed that cyanobacterial LPOR can oligomerize on membranes under certain conditions, forming stable oligomers with a mixture of a product and a substrate of the reaction, that is either Chlide and NADPH, or Pchlide and NADP+, on pure PG membranes (Fig. 1E-G, 6B). While pure PG membranes are not naturally present in Synechocystis cells, the observed oligomers may be an artifact of our in vitro system. Furthermore, the quantity of SynPOR in cyanobacterial cells is probably insufficient for effective oligomer formation, given the absence of reports on its accumulation, as observed in PLB. This could be attributed to the presence of dark-operative Pchlide reductase (DPOR)^33^, suggesting that cyanobacteria may not need to rapidly activate Chl synthesis in the light. However, the effective formation of SynPOR oligomers in vitro, especially with Pchlide and NADP+ supplementation, suggests a strong interaction between LPOR subunits promoted by PG, potentially useful for storing Chl intermediates during unfavorable conditions. This hypothesis, however, requires further experimental verification.

The presented data (Fig. 1) suggest that MGDG and PG have opposite effects on SynPOR, either promoting the release or the binding of dinucleotides, respectively. Such a regulatory role of lipids has not been reported for the cyanobacterial LPOR isoform yet, but it could play an important physiological role in vivo. By adjusting the relative MGDG:PG ratio, Synechocystis may regulate the rate of Chlide release from LPOR, directly impacting the rate of chlorophyll biosynthesis and potentially its cellular content. Studies have shown that in the Synechocystis mutant ΔpgsA, PG starvation is associated with decreased levels of Pchlide and increased levels of Chlide, among other alterations^34^. This observation aligns with the data presented in this study: the absence of PG cannot stabilize dinucleotide binding and the formation of Chlide:SynPOR:NADP(H) complexes, while MGDG promotes Chlide release, leaving SynPOR ready to convert more Pchlide molecules. We also predict that mutants lacking MGDG should exhibit increased Pchlide levels and decreased Chlide levels, as the absence of MGDG would disrupt products release from SynPOR, ultimately leaving all LPOR proteins occupied with the reaction products. However, such data has not been reported yet for cyanobacteria but *A. thaliana* MGDG deficient mutant lines were only able to germinate as small albinos growing heterotrophically ^35^.

### Evolutionary modifications towards modern plant LPOR

Based on a phylogenetic analysis of hundreds of LPOR sequences^4^, and considering the experimentally verified role of the residues at positions 122, 312, and 318 (Fig. 3), it can be deduced that gymnosperms and non-vascular plants have already developed modifications that render LPOR dependent on anionic membranes to effectively bind substrates (Fig. S7). Moreover, the sequences of the Pchlide loop and helix α10, that are conserved in all plant LPOR isoforms, have been already developed by algae (Fig. S10). These observations suggest that ancestral LPOR variant of land plants shared similarities in lipid-driven activity with AtPORA/B. While experimental verification of this claim is necessary, the data presented in this study, along with previous analyses^4^, indicate an intriguing evolutionary shift in LPOR properties between cyanobacteria and plants.

In cyanobacteria, LPOR activity appears to be independent of lipids, with membrane interaction primarily affecting product release post-reaction. In contrast, plant variants exhibit sequence alterations in helix α10 and the NADPH binding mechanism, rendering the enzyme effective in substrate binding only in the presence of anionic lipids, thus forming active complexes on the membrane. These adaptations enable plant LPORs, but not cyanobacterial variants, to efficiently store active complexes on membranes for extended periods. This evolutionary shift required several alterations in the enzyme sequence and was likely a gradual process.

We speculate that one of the first steps in this process was to involve the dependency of the substrate binding on the presence of anionic lipids (Fig. 6C). At the same time, the nature of the interaction with MGDG must have been changed. For plant LPORs, but not for the cyanobacterial homolog, the interaction with MGDG on the membrane results in a red-shift of the emission maximum of the complex^12^ and triggers LPOR oligomerization^13^, both of which significantly impact enzyme activity. The red-shift in emission maximum allows these complexes to potentially accept excitation energy transferred from other Pchlide:LPOR complexes, what improves efficiently of the reaction^36^. Additionally, MGDG-dependent oligomerization prevents accidental disassembly of active complexes, which is particularly beneficial for prolonged growth in darkness.

The sequence analysis revealed that the residues interacting with MGDG in AtPORB were already conserved in at least some algal LPOR sequences (Fig. S10E), suggesting that the process of LPOR alteration towards the plant-like variants started in algae. In contrast, the two oligomerization interfaces that are responsible for oligomerization of plant LPORs (Fig. S10C-F), surprisingly exhibit a diversity between cyanobacterial, algal and plant variants. Oligomerization interface I emerged as a small insertion in an ancestral algal LPOR sequence and was then modified in plants (Fig. S10C). One the other hand, oligomerization interface II emerged first as a substantial insertion in an ancestral homolog of algae, which was then eliminated in plants (Fig. S10D). This observation suggests that LPOR from algae may interact with MGDG similarly to plant variants, potentially leading to the formation of complexes with a characteristic emission maximum at 655 nm, although resulting oligomers are likely diverse, due to the variation in the oligomerization interfaces. It also elucidates the properties of a hypothetical second intermediate LPOR homolog, situated between cyanobacteria and plants. This enzyme likely underwent adaptations for prolonged interaction with the membrane in a form of the active ternary complex. At the same time, the oligomerization interfaces were extensive modified, probably to promote oligomerization and, coincidentally, to remodel membranes as a result of subunit interactions (Fig. 6D).

The open question remains what conditions were significant enough to select for LPOR homologs with such properties and when this selection occurred. The comparison of sequences suggests that the conditions favoring the selection of plant-like LPOR arose prior to the appearance of plants but after the endosymbiosis of chloroplasts, during the diversification of algae (Fig. S10). This is evident as certain algal sequences, like those from Chlamydomonas, closely resemble plant variants and are capable of forming PLB-like structures ^17^. Intriguingly, colonization of marine ecosystems by green algae (850-650 MYA) ^37^ coincides with global glaciations around 700 MYA ^38^. During this period lasting for tens of millions of years, called Snowball Earth event, ice sheets are believed to have covered nearly the entire surface of the planet. It was shown that low temperature and exceedingly dim light with sporadic events of light exposure influenced the genome of *Prochlorococcus*^39^, but certainly affected all the species, especially autotrophs. Any organisms capable of initiating photosynthetic activity the fastest during unpredictable light exposure events would have had a competitive advantage. We hypothesize that this was a driving force that shaped the evolution of LPOR towards plant-like variants during the Snowball Earth event.

### MV- and DV-Pchlide binding by LPOR

We have shown that neither LPOR from cyanobacteria nor from plants can discriminate between MV- and DV-Pchlide in a lipid free environment. Similar results for cyanobacterial and plant variants were obtained in previous studies: in lipid-free reaction mixtures SynPOR, LPOR from *T. elongatus* and AtPORC showed no preference towards MV- or DV-Pchlide^40,41^. Interestingly, variants from aerobic anoxygenic phototrophs exhibited a clear preference for either MV- or DV-Pchlide in lipid-free reaction mixtures ^40^, suggesting that these bacteria may have altered the pigment binding pocket of their LPOR variants.

Surprisingly, TaPOR2 ternary complexes with MV- and DV-Pchlide shown small but substantial difference in MGDG-induced oligomerization (Fig. 4A). The nature of the interaction between the group at C8 of Pchlide and the glycerol moiety of MGDG requires a dedicated quantum simulation study. Additionally, other plant LPOR isoforms should be tested in the presence of MGDG to determine if such a small difference in the tendency to oligomerization between MV- and DV-Pchlide can have any physiological effect. Nevertheless, even a slight acceleration of the onset of photosynthetic activity can be beneficial under challenging conditions and intense competition among photosynthetic species. We speculate an effecient formation of oligomers with MV-Pchlide comparing to DV-accelerates chlorophyll biosynthesis after a prolonged darkness. If MV-Pchlide is preferentially sequestered in LPOR oligomers, it is MV-Pchlide molecules that interact with MGDG in the binding pocket. This red-shifts their emission maximum makes them acceptors of absorbed excitation energy from complexes with DV-Pchlide. This implies that after exposure to light, MV-Chlide is formed first, which can be directly converted to Chl. Moreover, if MV-Pchlide is preferentially sequestered in LPOR oligomers, the remaining DV-Pchlide becomes available for DV-reductase. This enzyme can operate in darkness, converting DV-Pchlide to MV-Pchlide, thereby advancing Chl biosynthesis. This is a clear evolutionary advantage especially in the extreme environment of Snowball Earth. On the other hand, modern day DV-reductase of rice have been shown to accept DV-Pchlide, DV-Chlide and DV-Chl^42^, while the variant of Arabidopsis shown higher specificity towards DV-Chlide than DV-Pchlide^43^. These properties may be adaptations of the enzyme to the regular day and night periods characteristic of terrestrial environments.

### Prolamellar body formation

We have demonstrated that the addition of NADP+ and Pchlide induces the oligomerization of both plant LPORs and SynPOR, albeit on membranes with different composition. Moreover, the resulting complexes inherently differ in terms of shape and size. This could be an example of convergent evolution, suggesting that there is a universal physiological advantage to storing Pchlide when NADP+ concentration is high.

Regardless of the underlying factors leading to the ability to form PLB, the data presented in this paper suggest that LPOR is solely responsible for this process. The formation of the cubic phase as a result of NADP+ and Pchlide binding by LPOR in the presence of lipids containing MGDG indicates that it may be the initial stage of PLB formation (Fig. 6F). This also implies that the NADPH concentration must be low compared to NADP+, at least in the first days of seedling germination in darkness. If the NADPH concentration were high, leading to the direct formation of active ternary LPOR oligomers, the resulting complexes would be linear, not cubic (Fig. 6E). Based on our data, we can also predict that after cubic phase formation, the NADPH concentration within the etioplast should increase, which would result in dinucleotide displacement in the binding pocket of the enzyme and the formation of active ternary complexes (Fig. 6F). We have demonstrated that the formation of the complexes in this order on membranes preserves the cubic phase of the lipids with active ternary LPOR complexes (Fig. 5H, Fig. 6E).

However, the cubic phase obtained in our experiments is much less ordered than the ultrastructure of PLB. This suggests that the process of NADP^+^ displacement by NADPH is slow and gradual in vivo, allowing the slow crystallization of LPOR helical assemblies on the cubic lattice, what ultimately leads to the emergence of the characteristic paracrystalline structure. Perhaps, some other proteins may be involved in the formation of the characteristic pattern of PLB, but the involvement of LPOR in the formation of the cubic phase seems convincing. All the ligands required by the enzyme to remodel membranes into cubic phase are present in etioplasts, and no other process of cubic phase emergence have been reported so far that could explain the mechanism of PLB formation.

Based on our data, the molecular explanation behind of cubic phase formation by LPOR is not yet clear. A high-resolution structure of the enzyme within these complexes is required to fully address this question. Potentially, LPOR utilizes the property of MGDG, deforming the lipid membrane in such a way that the cubic phase becomes energetically favorable, owing to subunit interactions. That would explain why high salt concentrations and low pH deform PLB ultrastructure^23,44^ – by affecting the interactions between LPOR subunits that maintain the stability of the lipids in a phase that is not energetically stable without them. It also explains the mechanism of PLB disassembly after illumination^8^ – after the reaction, LPOR subunits of the oligomers dissociate from each other, allowing the membranes to relax and/or be modified by other proteins, such as CURT1^15^.

If the interactions between LPOR subunits that mainly are responsible for the formation of a cubic structure of PLB, the analysis of LPOR sequences provide an interesting view into the evolution of this mechanism. The diversity in the oligomerization interfaces in algal sequences (Fig. S10 CD) suggest that different orientation of subunits within the oligomers have been developed by these organisms. Out of them, at least one turned out to remodel lipids into the cubic phase in the presence of NADP^+^, which apparently was particularly advantageous, since all plants inherited this LPOR homolog. The ability to remodel membranes into cubic phase saturated with Chl intermediate in darkness is another adaptation of the enzyme that makes it optimized for prolonged storage and propel onset of photosynthetic activity. Storage is enhanced since the cubic phase allows for the accommodation of large bilayer areas, necessary for photosynthetic membranes, in a compact space. Lipid synthesis can occur in darkness, shortening the time needed to initiate photosynthetic activity and providing an evolutionary advantage. At the same time, the onset of photosynthetic activity is accelerated, as PLB serves as a reservoir of lipids and Chl intermediates that self-disassemble upon illumination, thereby triggering the system to commence under the appropriate conditions. Both of these adaptations seem to be particularly useful in the extreme environment of Snowball Earth, but also during the germination in highly competitive terrestrial environments. However, to accurately determine the origin of PLBs, a dedicated phylogenetic analysis of LPOR sequences in algae is required, supported by biochemical verification of their properties, including a structural investigation.

### LPOR variants in angiosperms

Hundreds of millions years after algae probably developed the mechanism of PLB formation, multiple duplication events in angiosperms resulted in the formation of second and third isoform of LPOR within this clade. Two main duplication events, occurring relatively recently, approximately 150-170 million years ago ^4^, led to the separate formation of two variants in monocots and eudicots. The presence of an additional copy of the gene enabled plants to explore the physiological advantages conferred by modified LPOR properties. At the time of the duplication events, the enzyme was already highly evolved and advanced. The sequence of helix α10 had been modified, and oligomerization interfaces had been perfected to produce tight helical arrangements. Moreover, the energy barrier between the open and closed active conformations was significantly elevated compared to cyanobacterial variants, necessitating substrate binding on the membrane. The former appeared to be easily reversible for newly formed spare copies of the LPOR gene, requiring just a handful of mutations near the critical residues responsible for NADPH binding. We identified three such candidates of amino acid residues at positions 122, 312, 318, which are in close proximity to three highly conserved positively charged residues that interact with the phosphate moiety of NADPH, namely R121, K125, and R317. The residues 122 and 318 likely utilize a similar mechanism to alter the size of the barrier between open and closed state. If a negatively charged residue is present at either of these positions, it competes with the phosphate group of NADPH for interaction with the positively charged residues at 121, 125, and 317. Substituting a negatively charged residue with a neutral one eliminates this competition, promoting a conformational change towards the closed conformation. On the other hand, the residue at position 312 is close to a positively charged K332 of helix α10. The introduction of a negatively charged residue at position 312 enables the formation of a strong salt bridge, stabilizing the closed conformation of the helix α10.

In clade C, we identified three different combinations of residues at positions 122, 312, and 318 that, according to the measurements, reduce the energy barrier between the open and closed states, allowing the enzyme to function effectively without the aid of anionic membranes. Meanwhile, in clade AB, the combination of residues at positions 122, 312, and 318 is highly conserved not only within the clade but also in monocots. According to measurements, this set of residues, namely D, T, and E, respectively, makes the conformational change more difficult in a lipid-free environment, but their presence overcomes this barrier in the presence of anionic lipids. We can speculate that all isoforms sharing identical residues at positions 122, 312, and 318 with AtPORA, AtPORB, and the monocot isoforms rely on anionic lipids for substrate binding. However, no predictions can be made about the ability to form cubic phases from this, as these residues are not involved in the interactions between LPOR subunits.

The monocot isoforms, although sharing identical residues at positions 122, 312, and 318, exhibit differences in activity when no lipids are added to the reaction mixture, with TaPOR1 requiring much higher NADPH concentrations to achieve the same activity as TaPOR2 (Fig. 3D, Fig.S7). This observation suggests that other positions may influence the energy barrier between the open and closed states of LPOR. Based on sequence comparisons, we suggest that the residue at position 178 could contribute to the observed difference in activity. It is close to the residue involved in NADPH binding, namely E318, and potentially can interfere with the conformational change of the enzyme. All M1 variants have a conservative R residue at this position, while in other clades, it is either a Q, F, or L residue. The mechanism by which a positively charged residue at position 178 could hinder active complex formation requires experimental verification.

Nevertheless, based on the data obtained so far, it seems plausible that both monocots and eudicots independently developed two isoforms of LPOR starting from an ancestral homolog that works preferentially at lipid membranes. Eudicots modified their novel isoform so it can bind substrates in a soluble form, while monocots made a newly obtained LPOR copy even more dependent on the lipids. It must be emphasized that anionic-lipid-assisted NADPH binding is just one aspect in which variants can differ. Studies have shown that LPOR variants of barley vary in Pchlide binding affinity^45^, while other variants differ in optimal pH and/or temperature regarding their activity^40^. Certainly, more variants of angiosperms need to be characterized to identify differences not only in substrate binding and activity, but also in product release, the ability to form cubic phase, and the strength of interaction between LPOR subunit within oligomers. Such characterization may allow to suggest residues responsible for these properties, which would be useful for future site-directed mutagenesis studies and phylogenetic investigations.

## Materials and Methods

### Gene cloning

In this paper we used AtPORA_WT, AtPORB_WT, AtPORC_WT and SynPOR that have been cloned for a previous study^4^. *TaPOR1* gene and *TaPOR2* gene (Uniprot: Q41578, W4ZSC8) were cloned from Triticum aestivum seedling, using RNA extraction kit (Sigma-Aldrich) followed by cDNA synthesis with oligoT primers (ProtoScript II, New England Biolabs). The *TaPOR2* gene lacking the part coding a transit peptide and pET15b plasmid (Novagen) were amplified with PCR using Q5 polymerase (New England Biolabs) according to the manufacturer’s protocol: 98°C for 30 seconds; 30 cycles of 98°C for 10 seconds, 71°C for 30 seconds, 72°C for 40 seconds; final extension 72°C for 2 minutes, with specific primers (Table S2) The products of the reactions were purified from agarose gel using Gel Extraction Minipreps Kit (Bio Basic Canada) and ligated together with NEBuilder HiFi DNA Assembly Kit (New England Biolabs). Escherichia coli DH5α competent cells were transformed with the ligation mixtures and the cells containing assembled plasmid were selected on an agar medium supplemented with 100 mg/l ampicillin. The clones containing inserts were identified using colony PCR. The recombinant plasmids were isolated using plasmid purification kit (Bio Basic Canada) and the inserts were verified by sequencing (Genomed, Poland)

### Site-directed mutagenesis and hybrid variants

Point mutations were introduced into pET15b_AtPORB and pET15b_AtPORC plasmids using SLIM method ^46^ with specific primers (Table S2) as described in previous study^12^.

To produce a hybrid protein of AtPORB and SynPOR, a whole vector pET15B_AtPORB was amplified except for a part of gene that was supposed to be changed for SynPOR sequence. At the same time, appropriate fragment of SynPOR gene was amplified with specific primers with additional fragments corresponding to amplified pET15b_AtPORB (Table S2) with Q5 polymerase (New England Biolabs) according to the manufacturer’s protocol. Fragments were ligated together with NEBuilder HiFi DNA Assembly Kit (New England Biolabs). Then Escherichia coli DH5α competent cells were transformed with the ligation mixtures and the cells containing assembled plasmid were selected on an agar medium supplemented with 100 mg/l ampicillin. The clones containing inserts were identified using colony PCR. The recombinant plasmids were isolated using plasmid purification kit (Bio Basic Canada) and the inserts were verified by sequencing (Genomed, Poland).

### Protein expression and purification

All proteins were expressed and purified according to the previously published protocol with slight modifications^47^. Cultures of E. coli BL21(DE3)pRIL transformed with expression vectors were induced with 0.5 mM IPTG after OD reached 0.6 and incubated for 18h at 22°C with rotary shaking. In comparison to the previous protocol, the higher IPTG concentration and prolonged expression time allowed us to increase a yield of protein purification.

### Pchlide and Chlide purification

Pchlide was purified from etiolated wheat seedlings according to previous study^12^. Chlide was prepared from chlorophyll a according to the protocol^48^.

### Sample preparation and fluorescence measurements

Sample preparation was preformed according to the previous reports^12^. All reaction mixtures contained 37 mM sodium phosphate buffer (Na2HPO4/NaH2PO4; pH 7.1), 225 mM NaCl, 150 mM imidazole, 5 mM β-mercaptoethanol and 25% glycerol. The exact concentrations of the reagents are given in the descriptions of the figures. The range of the concentrations used in the experiments was 62.5 nM–200 μM for NADPH/NADP+ (AppliChem), 5 mM for Pchlide/Chlide, 15 μM for LPORs and 2–400 μM for the lipids (Avanti Polar Lipids). OPT lipids refer to the mixture of lipids optimized for previous structural study and contained: 50 mol% MGDG, 35 mol% DGDG and 15 mol% PG (Avanti Polar Lipids).

Samples of the proteins were stored at −20°C and were gently thawed on ice just before the preparation of reaction mixtures. The addition of the pigments to the reaction mixture was performed under a dim green light. The samples were incubated in darkness for 30 min at room temperature before the measurement, then placed in quartz capillaries and frozen in liquid nitrogen for fluorescence measurement at −196°C. After the measurements, the samples were thawed at room temperature under a dim green light and illuminated for 20 seconds or 20 minutes with white light of intensity 80 μmol photons m^−2^ s^−1^, then frozen again and measured again.

The low-temperature fluorescence spectra were measured with PerkinElmer LS-50B spectrofluorometer equipped with a sample holder cooled with liquid nitrogen The spectra were recorded between 600 and 790 nm with a scanning speed of 400-500 nm/min. The data collection frequency was 0.5 nm and the excitation wavelength was 440 nm. Excitation and emission slits were in a range 7-10 nm.

### Calculations for activity determination

The samples were prepared and illuminated for 20 seconds as described above. From the spectrum, the emission intensities of Chlide (F Chlide) and Pchlide (F Pchlide) were recorded at the peaks’ maxima: between 674-695 and 632-655, respectively (Fig. 2G). The ratio of F Chlide / F Pchlide was then calculated for each sample. For every analyzed protein, a modified Michaelis-Menten equation was fitted to the data: (v_max_ * [NADPH]) / (K_M_ * [NADPH]) + a. The modification lies in the “+ a” term, as for inactive samples, the ratio F Chlide / F Pchlide never reaches zero, given that Pchlide emits some light at 674-695nm (Fig. 2GH).

### Electron microscopy

Samples for electron microscopy were negatively stained according to the previously published protocol ^13^ and visualized with JEOL JEM2100 HT CRYO LaB6 electron microscope. At least three independent sample preparations were analyzed, and representative micrographs are presented. Samples for cross-section analysis of were prepared according to the previously described method^49^.

## Acknowledgments

We thank Jerzy Kruk for his help with MV- and DV-protochlorophyllide purification and Leszek Fiedor for providing a sample of purified chlorophyllide. This work was supported by SONATA project (2019/35/D/NZ1/00295) granted by NCN to MG. Part of the results presented in Fig. 2 were obtained thanks to the support form Faculty of Biochemistry, Biophysics and Biotechnology (N19/MNS/000026) granted to MG. The animation of chlorophyll biosynthesis was funded by the Priority Research Area BioS under the program Excellence Initiative – Research University at the Jagiellonian University in Krakow (B.1.7.2021.16) granted to MG. We thank Bernhard Grimm for helpful suggestions regarding the text of the manuscript.

## Authors contribution

WO: Investigation. KS: Investigation. MŁ: Investigation. OB: Investigation. MG: Conceptualization, Methodology, Validation, Writing - Original Draft, Writing - Review & Editing, Visualization, Supervision, Investigation, Funding acquisition.

## Supplementary materials

**Figure S1.**
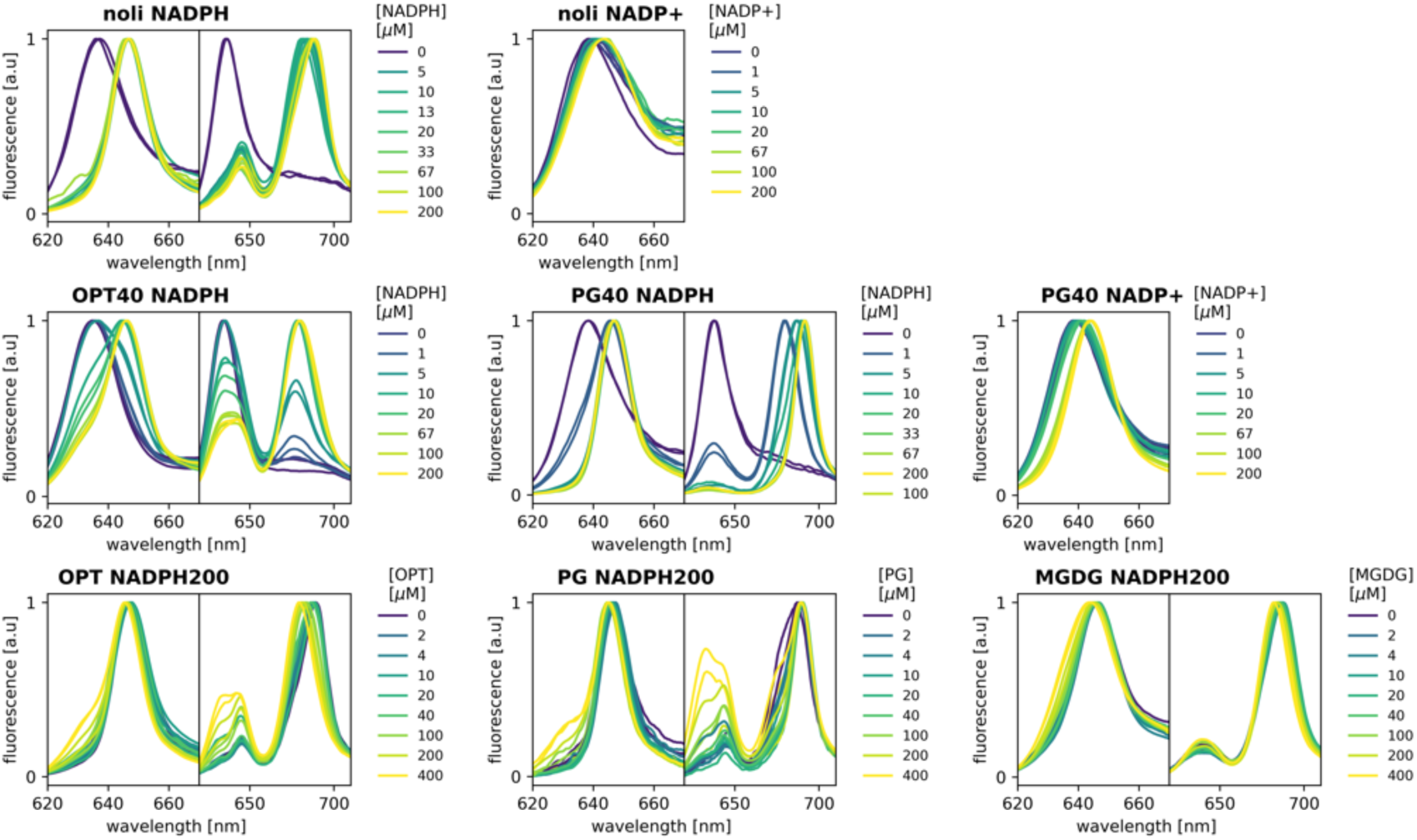
The spectra of SynPOR reaction mixtures used for Fig 1BD. The left panels present spectra before illumination, and the right panels present spectra after 20 seconds of illumination. “Noli” indicates no lipids added to the reaction mixture, “OPT/PG 40” indicates 40 µM lipids, and “NADPH200” indicates 200 µM NADPH.

**Figure S2.**
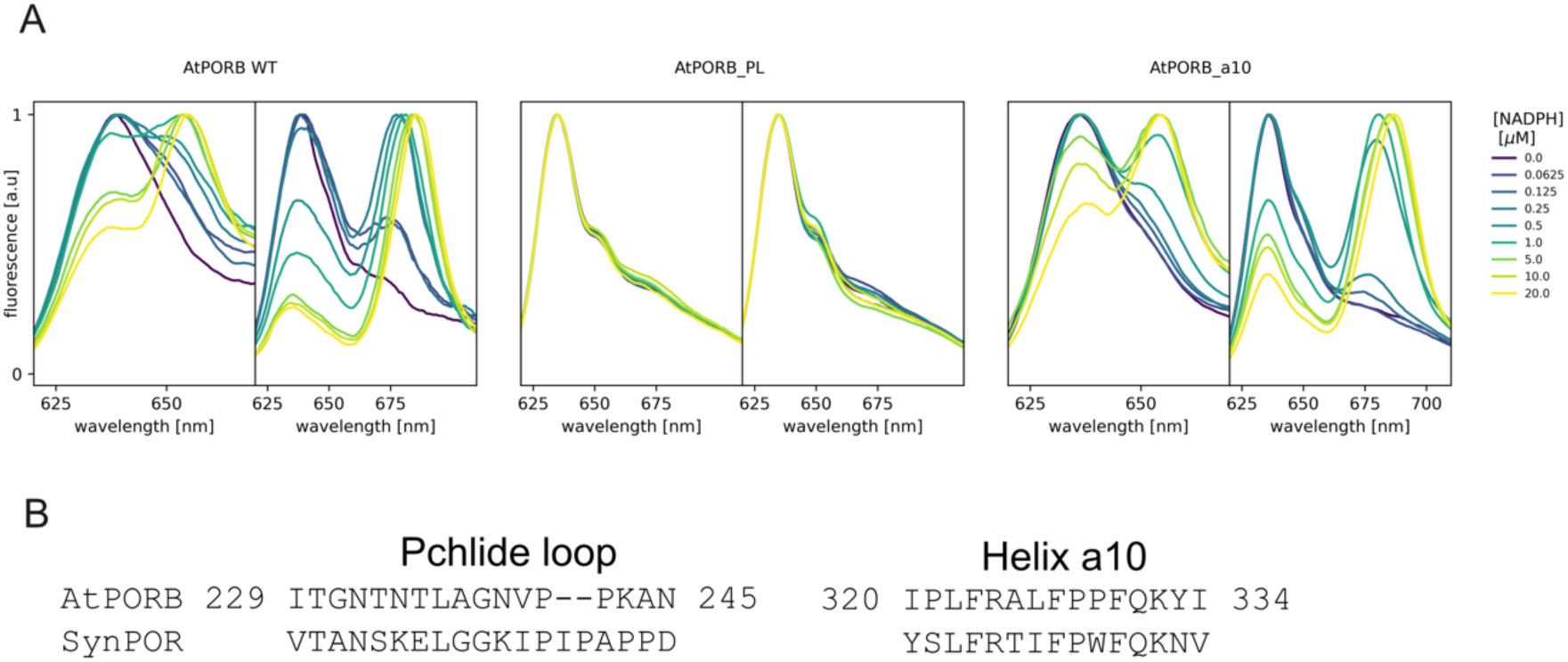
Hybrid LPOR variants. **A.** The spectra before (left panels) and after 20 seconds of illumination (right panels) for reaction mixtures of AtPORB WT and the hybrid variants in which parts of AtPORB were changed into the corresponding sequences from SynPOR. For AtPORB_PL and AtPORB_α10, the Pchlide loop and helix α10 were changed, respectively. **B.** The alignment of fragments of sequences corresponding to the Pchlide loop and helix α10 in AtPORB and SynPOR. The provided numeration of residues corresponds to the AtPORB sequence.

**Figure S3.**
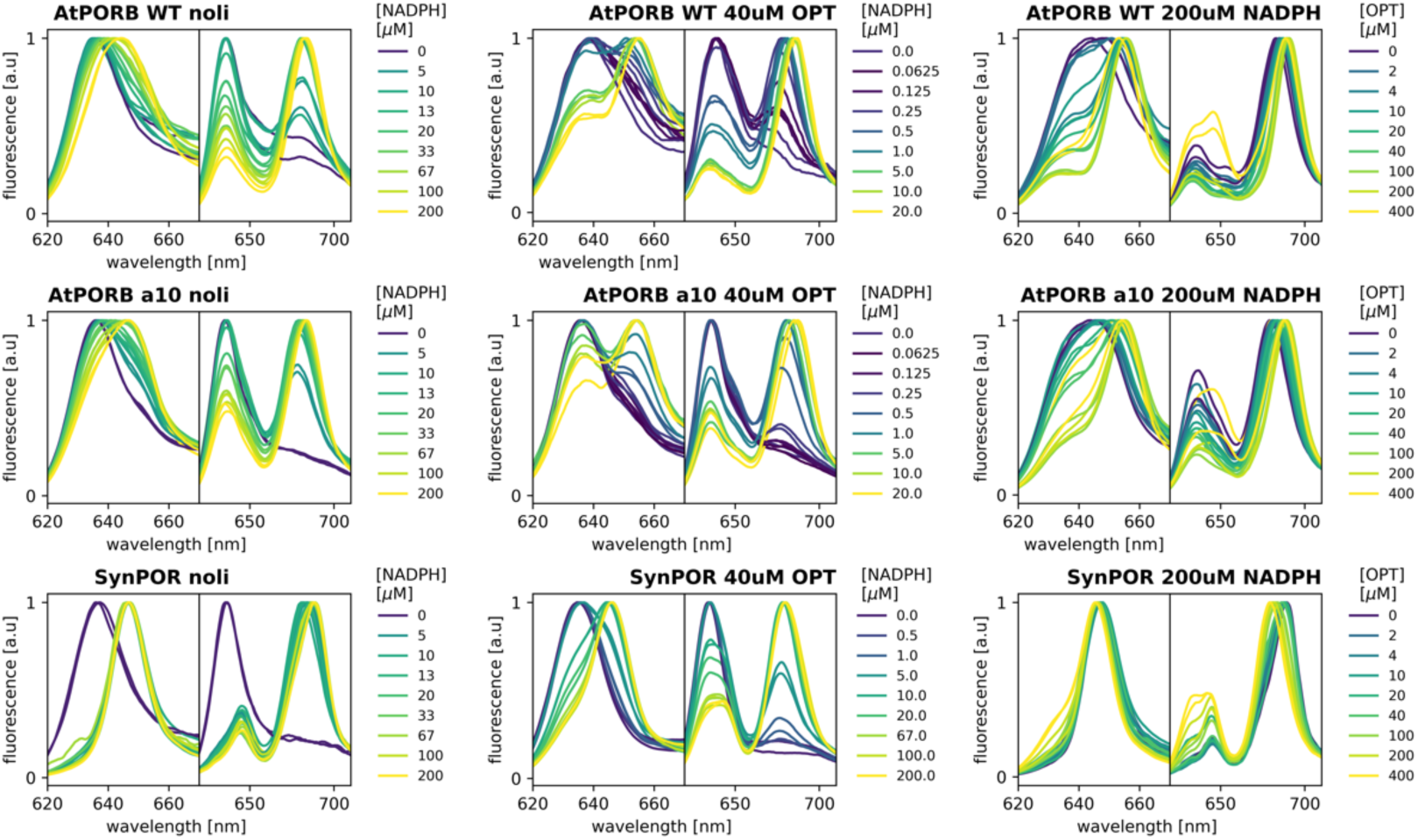
The spectra of SynPOR, AtPORB, and AtPORB_a10 reaction mixtures used for Fig 2CF. The left panels present spectra before illumination, and the right panels present spectra after 20 seconds of illumination. “Noli” indicates no lipids added to the reaction mixture, “OPT/PG 40” indicates 40 µM lipids, and “NADPH200” indicates 200 µM NADPH.

**Fig. S4.**
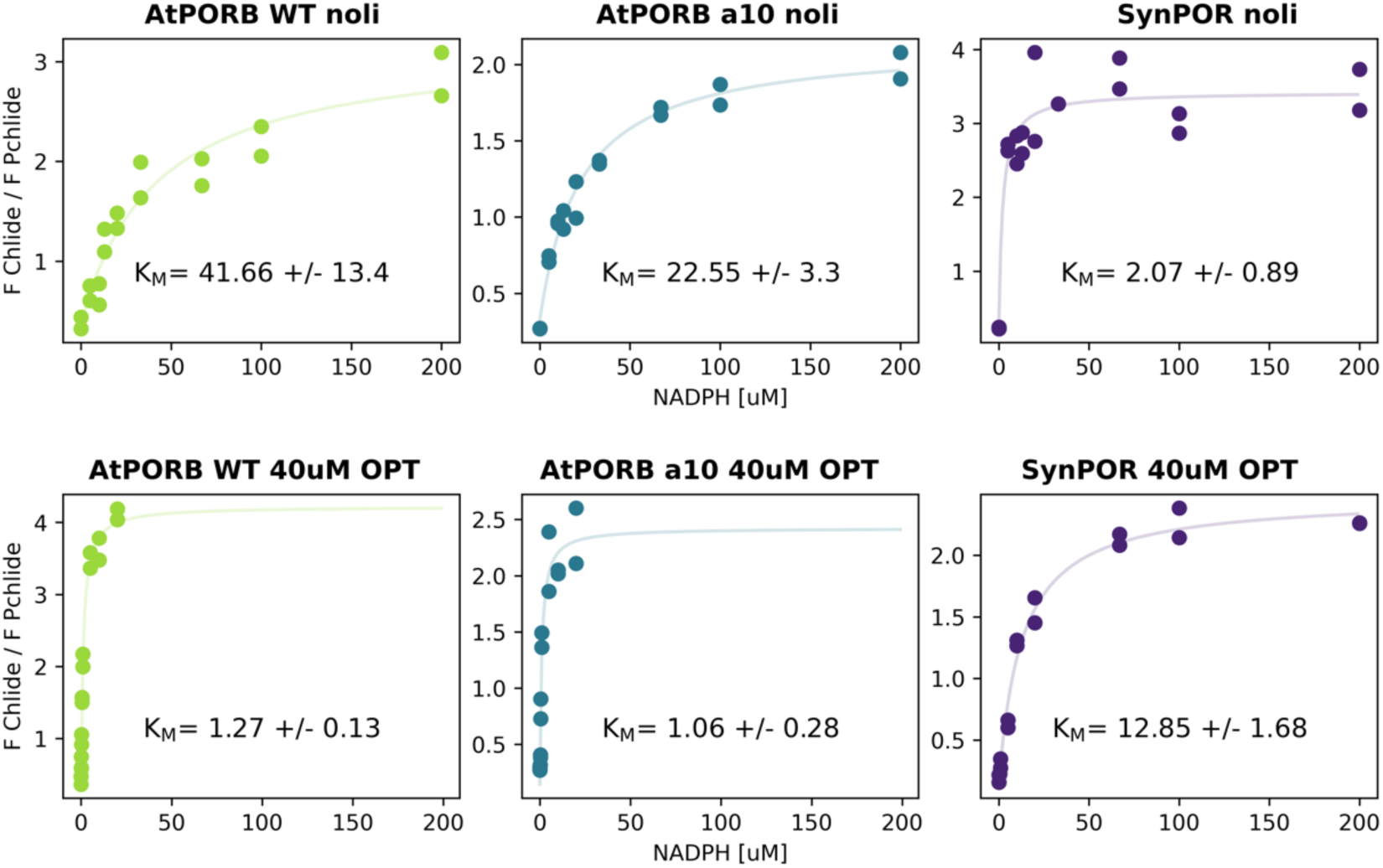
K_M_ determination for AtPORB WT, AtPORB_α10, and SynPOR. The relationship between F Chlide / F Pchlide and NADPH concentration in the absence of lipids (top panels) and with 40 µM OPT lipids (bottom panels). A fit of a modified Michaelis-Menten equation is shown with K_M_ value.

**Fig. S5.**
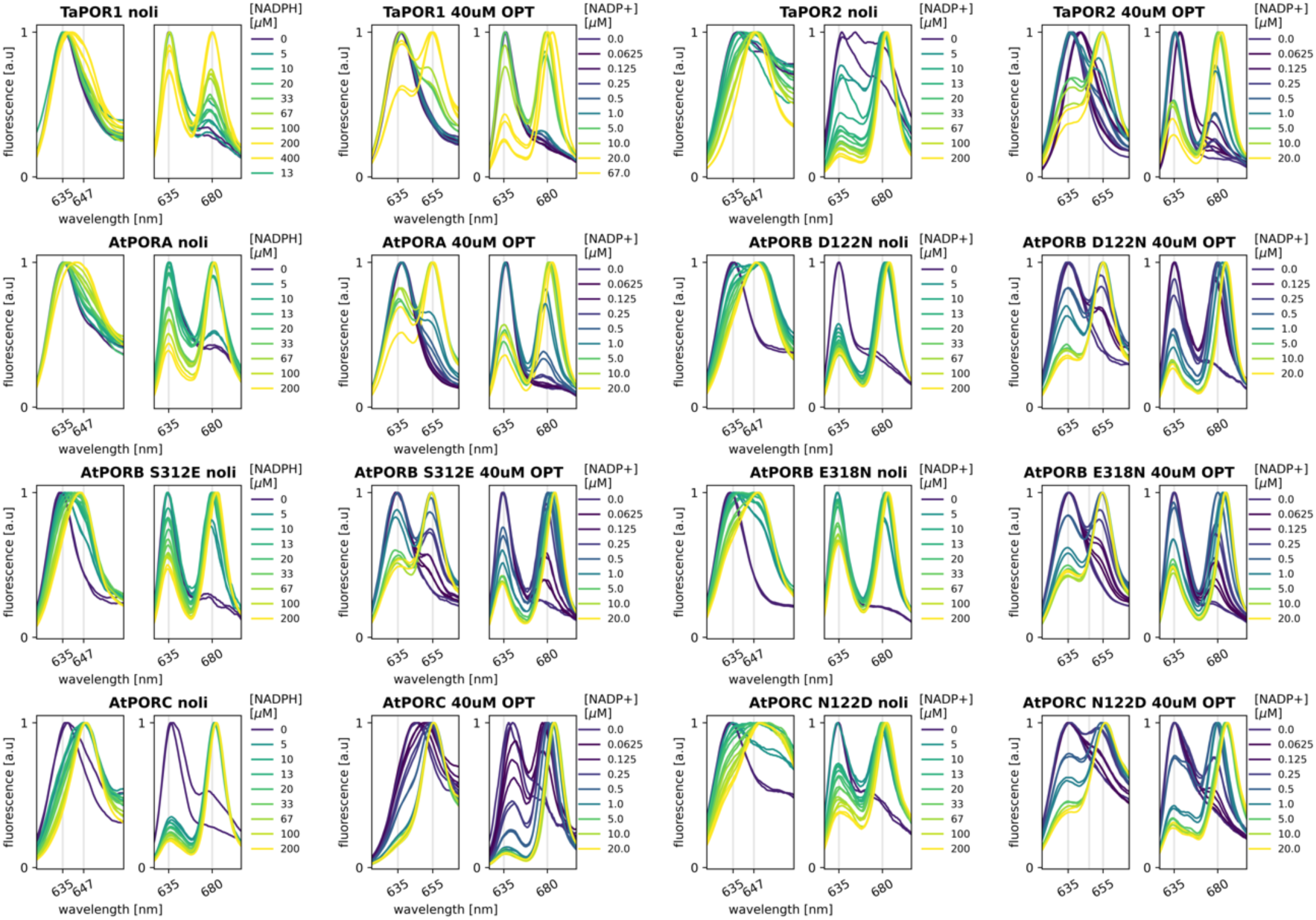
The spectra of reaction mixtures of WT LPORs and the mutants used for K_M_ determination. The left panels present spectra before illumination, and the right panels present spectra after 20 seconds of illumination. “Noli” indicates no lipids added to the reaction mixture, “ 40 uM OPT” indicates 40 µM lipids.

**Fig. S6.**
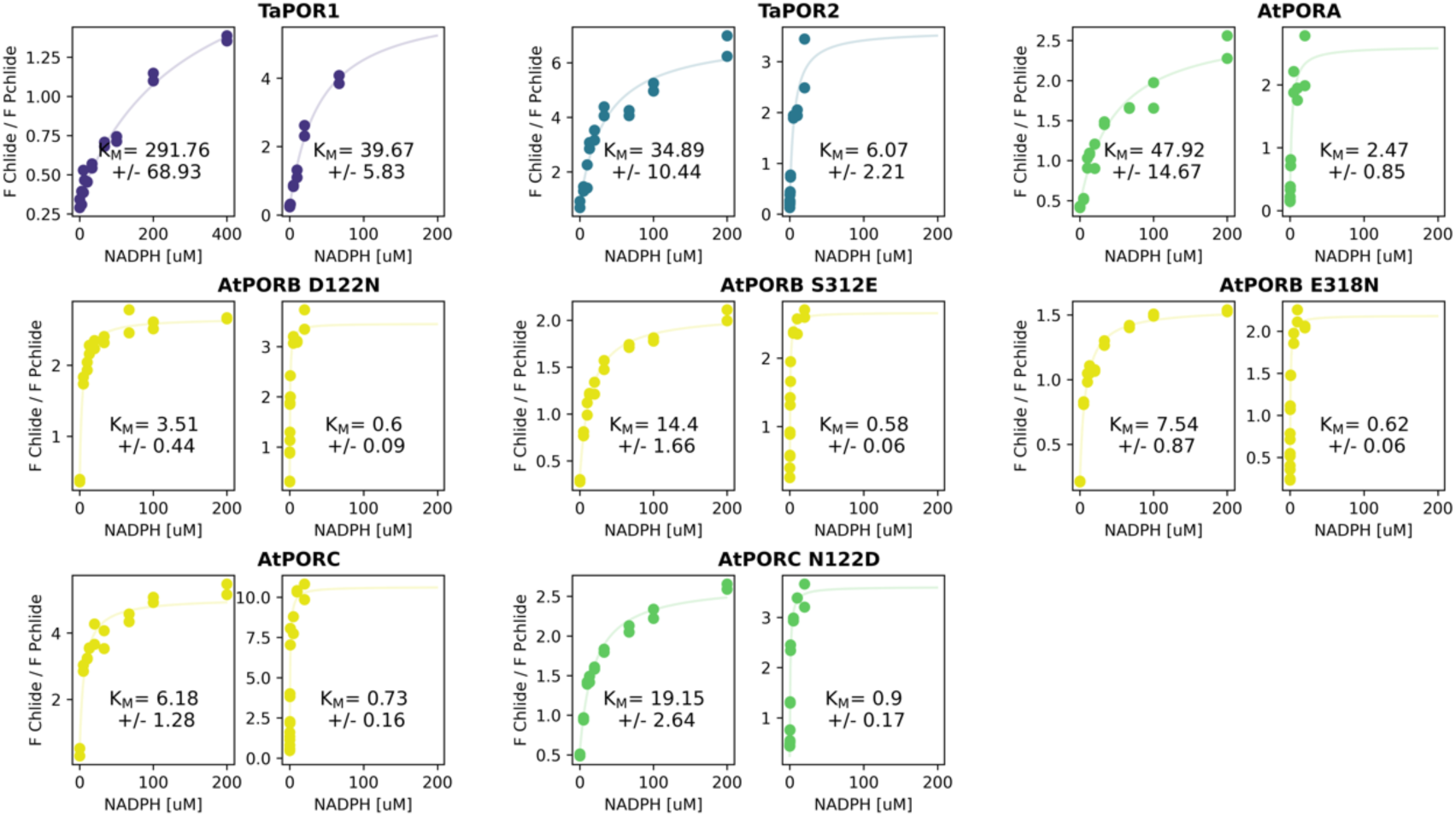
K_M_ determination for WT LPORs and the mutants. The relationship between F Chlide / F Pchlide and NADPH concentration in the absence of lipids (left panels) and with 40 µM OPT lipids (right panels). A fit of a modified Michaelis-Menten equation is shown with K_M_ value. Data for PORB WT are presented in Fig. S5.

**Figure S7.**
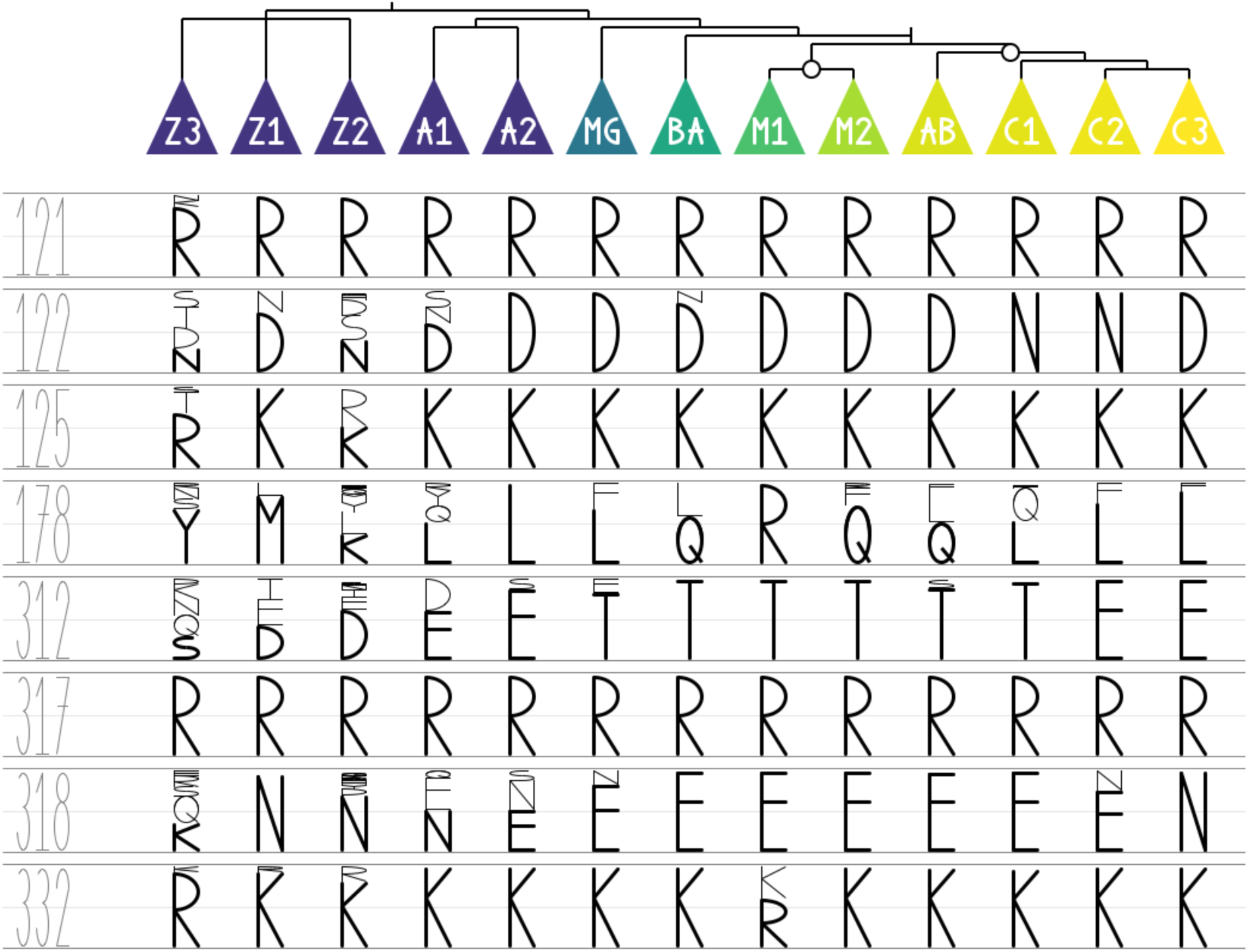
Conservation of selected positions in LPOR sequences. A simplified cladogram of LPORs showing evolutionary relationships between the clades and conservativeness of selected positions (numbering for AtPORB). Z3, Z1, Z2 – cyanobacterial clades (15, 176, 155 sequences, respectively); A1, A2 – algal clades Al1 and Al2 (10 and 9 sequences, respectively); MG – lower plants and gymnosperms (6 sequences); BA – basal angiosperms (7 sequences); M1, M2 – two clades of monocots (15 and 18 sequences, respectively); AB, C1-3 – clades of eudicots (81, 35, 4 and 11 sequences, respectively). The size of letters is scaled proportionally to the number of sequences that have a given amino acid residue at the given position.

**Fig. S8.**
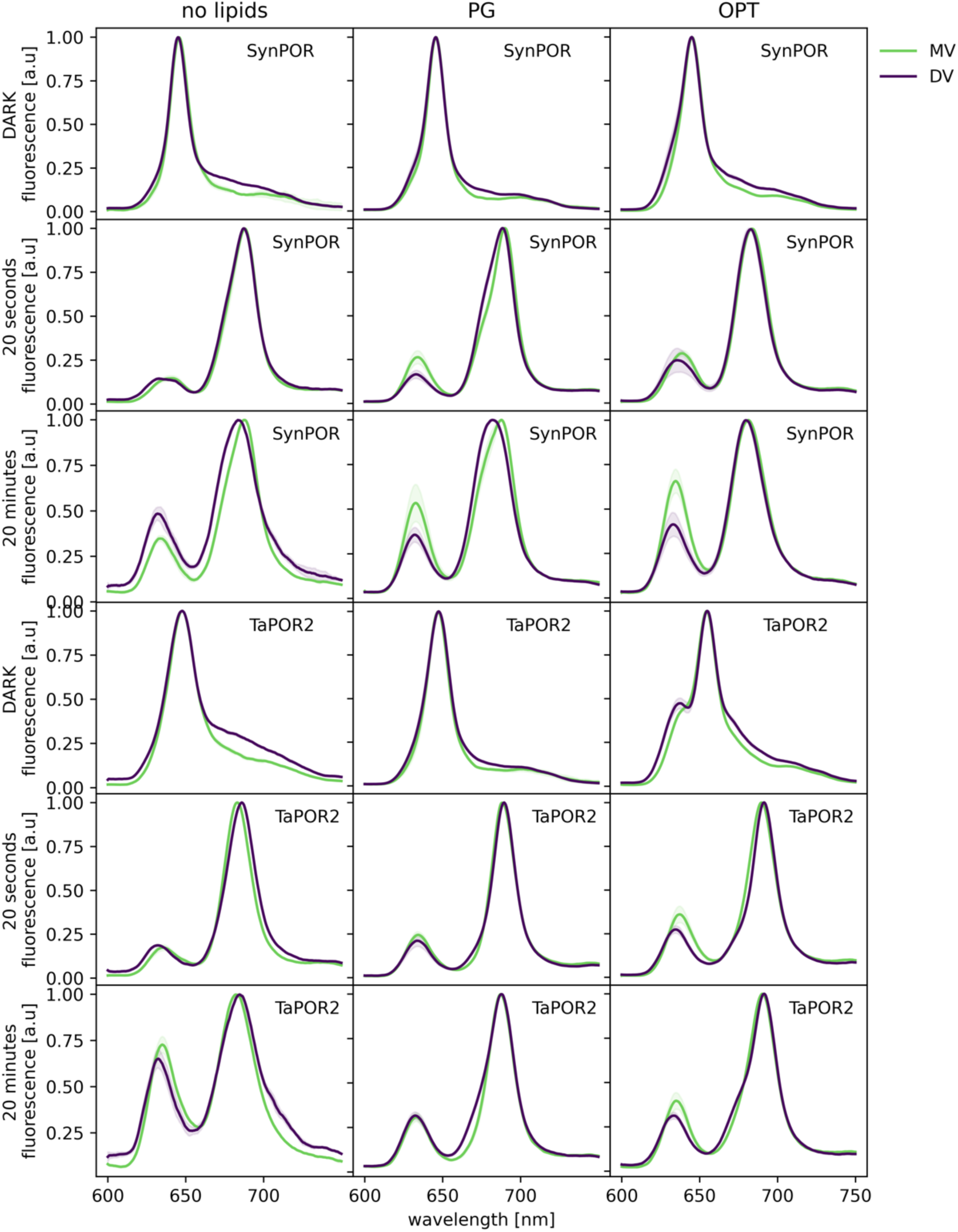
The spectra of MV- and DV-Pchlide in the reaction mixture containing SynPOR or TaPOR2. The reaction mixtures contained 15 µM LPOR, 3.2 µM Pchlide (either MV- or DV-), 200 µM NADPH, and either 100 µM PG, 100 µM OPT, or no lipids (Noli). The spectra were measured before illumination, after 20 seconds or 20 minutes of illumination (see y axis label). Shared areas represent the standard deviation between replicates.

**Fig. S9.**
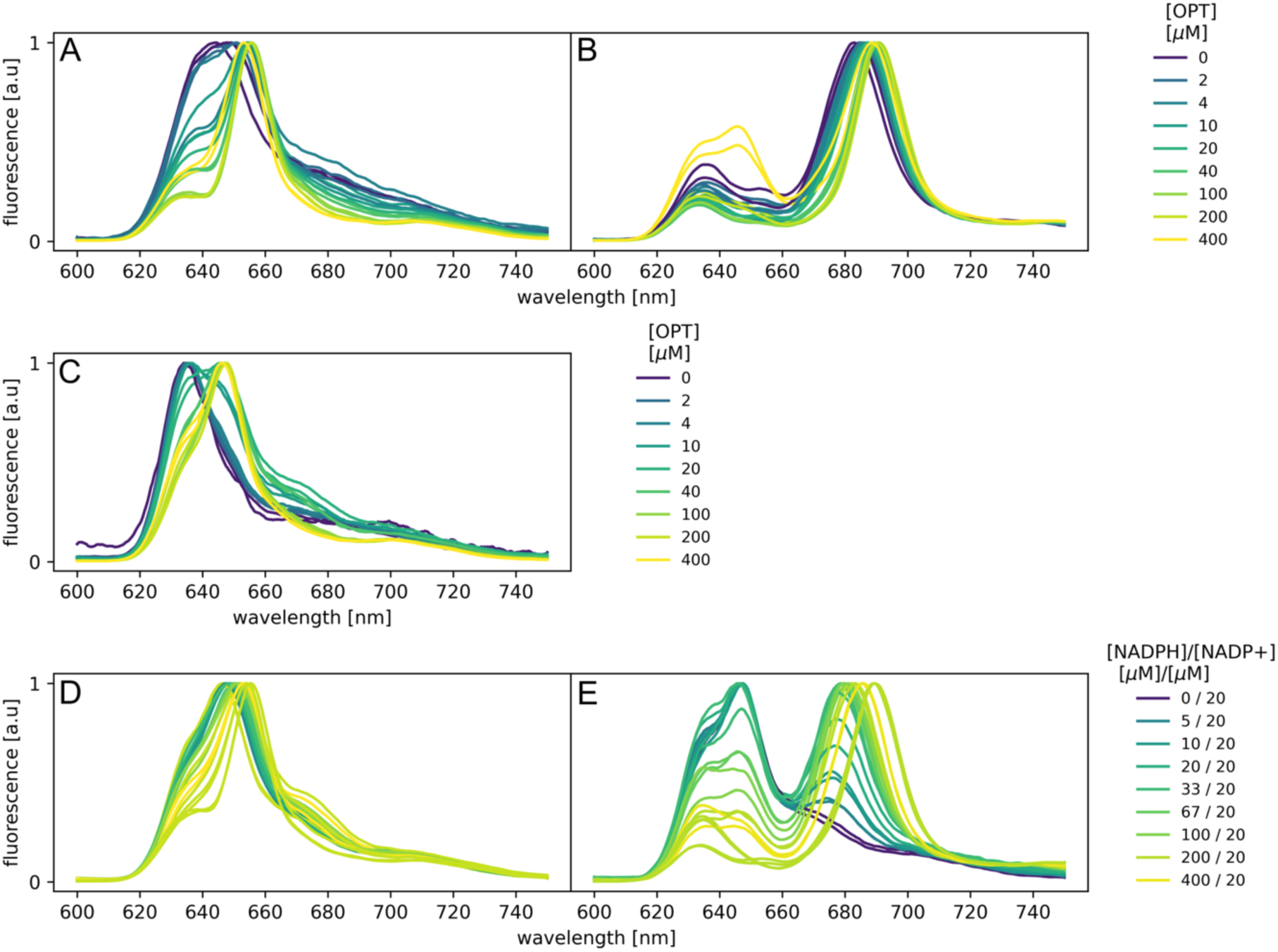
The spectra of the reaction mixtures of AtPORB with NADP+, NADPH or mixture of the two. AC. The spectra before illumination of AtPORB with 200 µM NADPH (A) or NADP+ (C). B. The spectra of samples presented in A illuminated for 20 seconds. DE. The spectra before (D) and after 20 seconds of illumination (E) of the reaction mixture containing a mixture of NADPH and NADP+ in the presence of µM OPT lipids. The concentration of ingredients: 5 µM Pchlide, 15 µM AtPORB.

**Figure S10.**
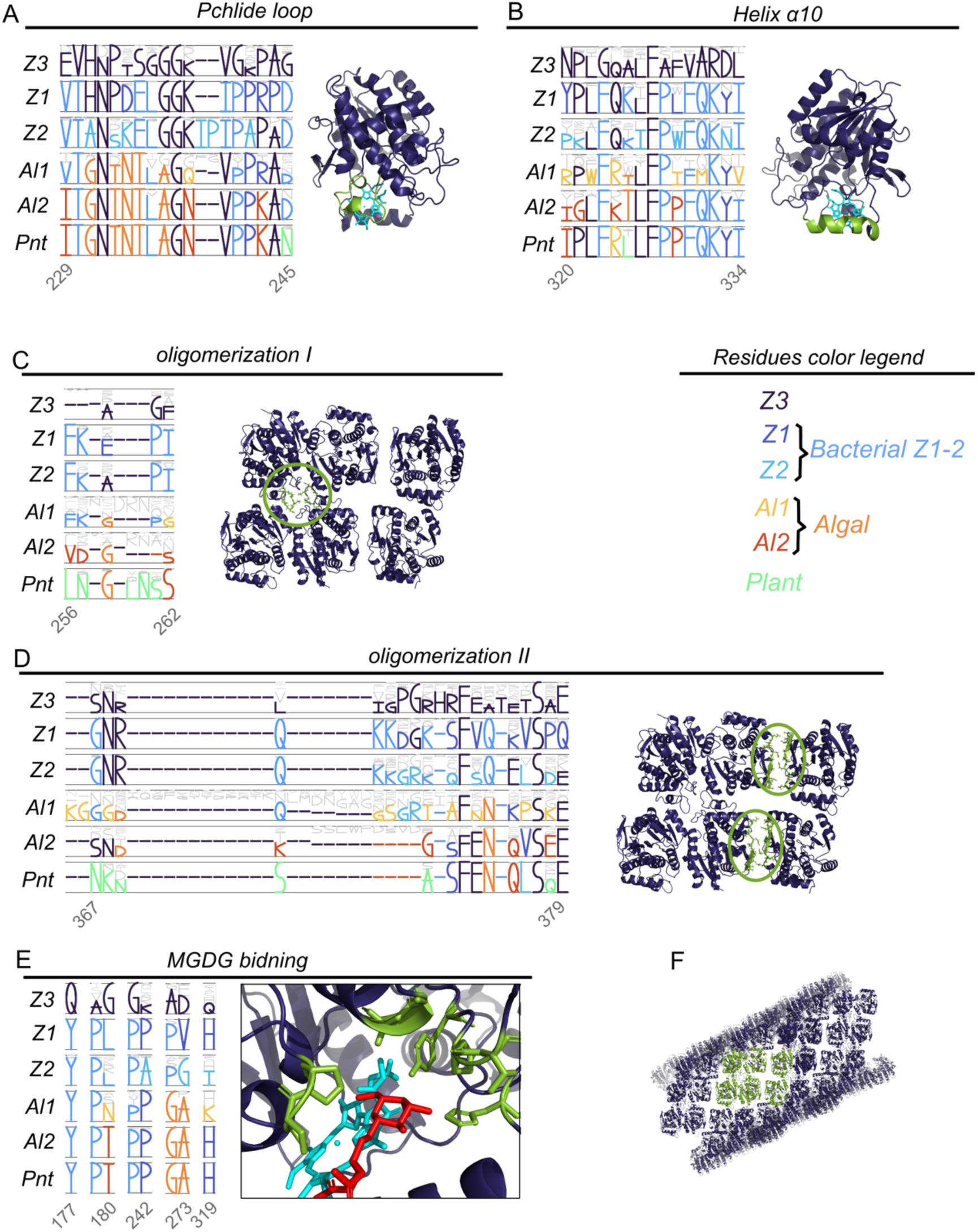
Conservativeness of selected elements in LPOR sequences. Z3, Z1, Z2 – cyanobacterial clades (15, 176, 155 sequences, respectively); Al1, Al2 – algal clades (10 and 9 sequences, respectively); Pnt – lower plants, gymnosperms and angiosperms (177 sequences). The size of letters is scaled proportionally to the number of sequences that have a given amino acid at the given position. The residues are color-coded according to the clade in which a given residue at a specific position became conservative for the first time. The corresponding parts of the sequence are marked on LPOR structure (pdb: 9JK7). For CD, the oligomerization interfaces are additionally marked with green ellipses. Fragment of the LPOR filament with six subunits, corresponding to those displayed in panels CD, is marked in green.

**Video S1.** Simplified but scientifically accurate visualization of chlorophyll biosynthesis reactions in plants.

**Table S1.**
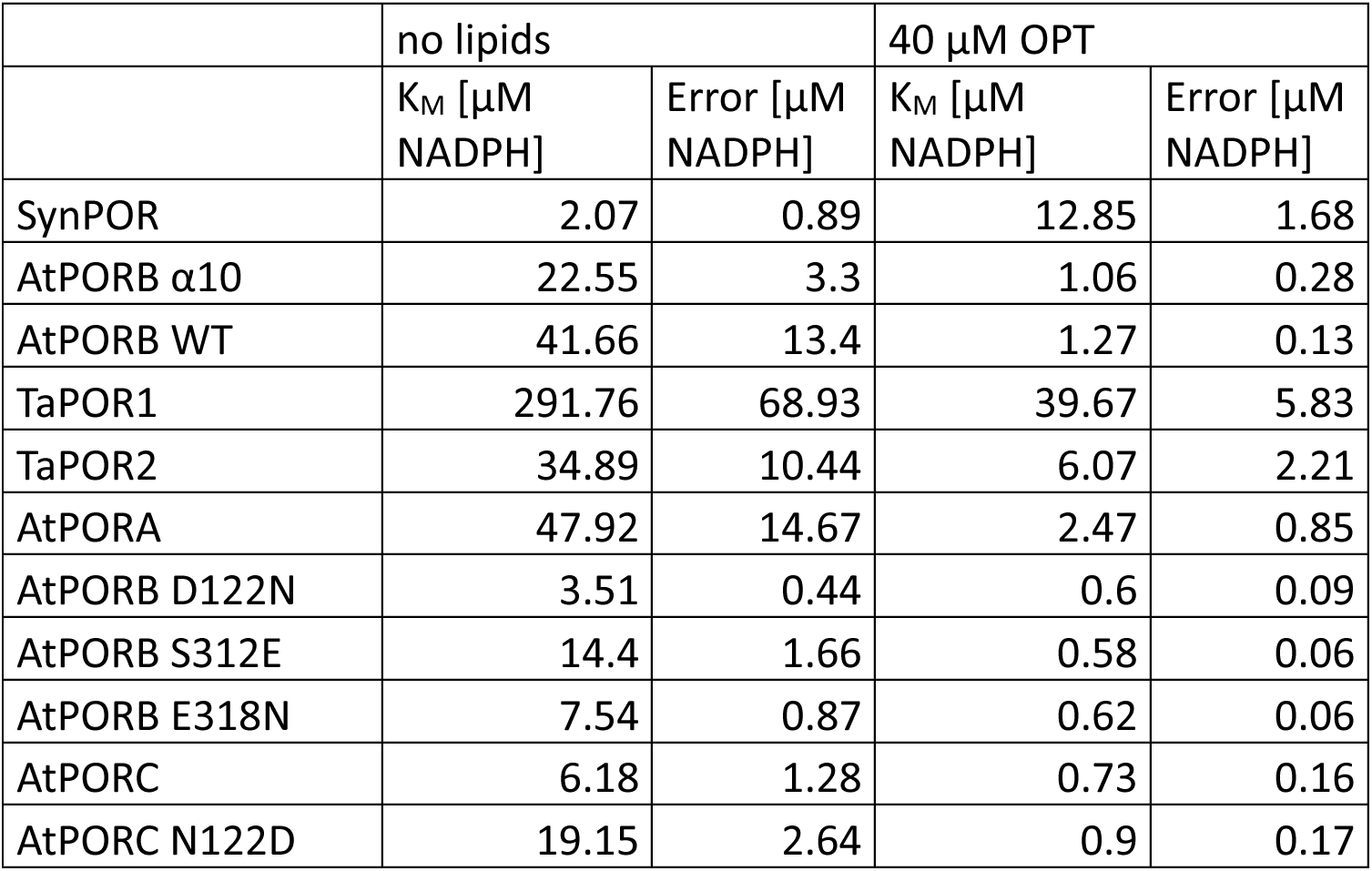
K_M_ values determined for WT LPORs and mutants in the presence and absence of 40 µM OPT lipids.

| | no lipids | | 40 $\mu\text{M}$ OPT | |
| --- | --- | --- | --- | --- |
| | $K_M$ [$\mu\text{M}$ NADPH] | Error [$\mu\text{M}$ NADPH] | $K_M$ [$\mu\text{M}$ NADPH] | Error [$\mu\text{M}$ NADPH] |
| SynPOR | 2.07 | 0.89 | 12.85 | 1.68 |
| AtPORB $\alpha 10$ | 22.55 | 3.3 | 1.06 | 0.28 |
| AtPORB WT | 41.66 | 13.4 | 1.27 | 0.13 |
| TaPOR1 | 291.76 | 68.93 | 39.67 | 5.83 |
| TaPOR2 | 34.89 | 10.44 | 6.07 | 2.21 |
| AtPORA | 47.92 | 14.67 | 2.47 | 0.85 |
| AtPORB D122N | 3.51 | 0.44 | 0.6 | 0.09 |
| AtPORB S312E | 14.4 | 1.66 | 0.58 | 0.06 |
| AtPORB E318N | 7.54 | 0.87 | 0.62 | 0.06 |
| AtPORC | 6.18 | 1.28 | 0.73 | 0.16 |
| AtPORC N122D | 19.15 | 2.64 | 0.9 | 0.17 |

**Table S2.**
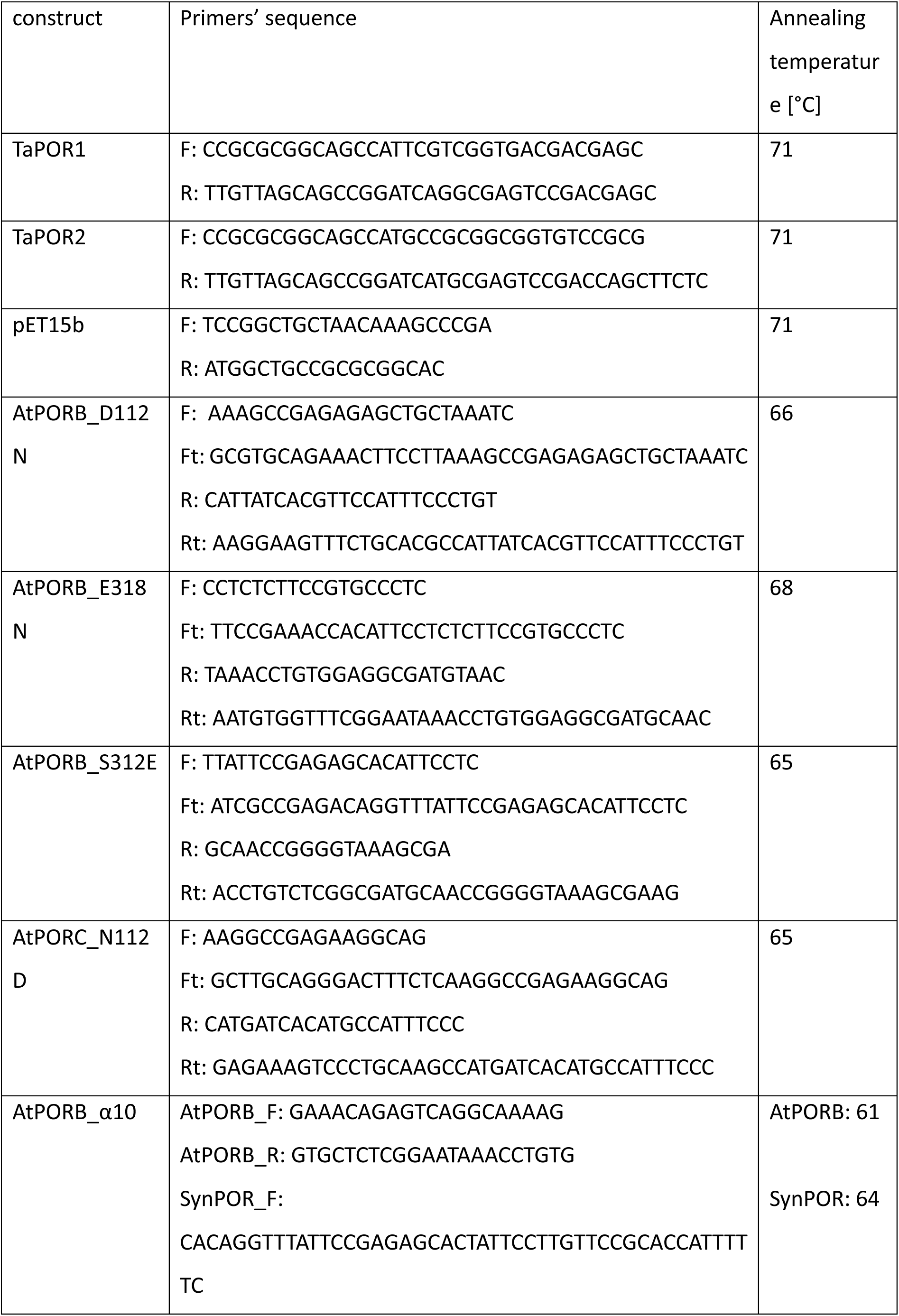
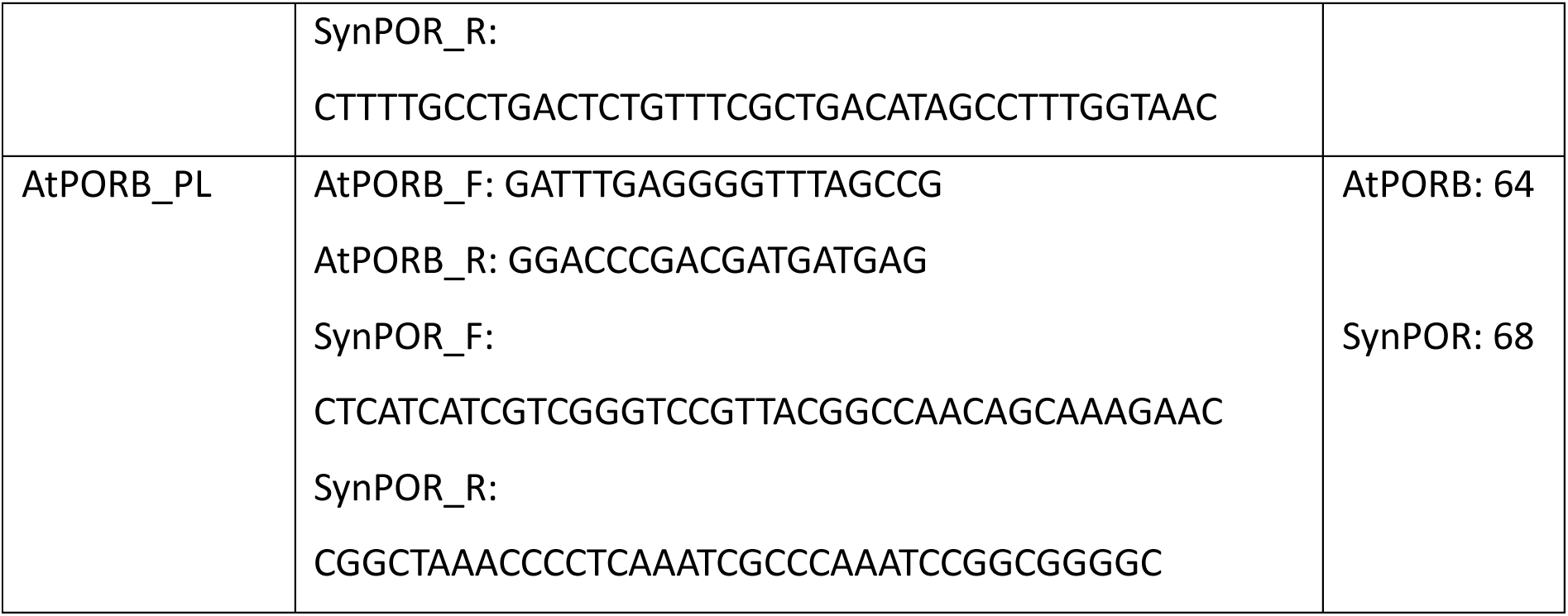
List of primers used in the study with annealing temperatures used.

